# Cooperative processing of visual threat across collicular pathways

**DOI:** 10.64898/2026.10.12.758428

**Authors:** Bram Nuttin, Arnau Sans-Dublanc, Anna Chrzanowska, Chen Liu, Zoi Drakaki, Laura Hanson, Rik Luykx, Amber Bruynsteen, Adrien Philippon, Micheline Grillet, Norma Kühn, Pedro J. Gonçalves, Gabriel Montaldo, Vincent Bonin, Alan Urban, Karl Farrow

## Abstract

Rapid defensive actions require the transformation of sensory cues into appropriate motor responses. The superior colliculus is central to this transformation, containing defined cell-types that form parallel output pathways linking visual inputs to distinct behaviors. However, whether these pathways act independently or cooperate remains unclear. Here we combine cell-type-specific inhibition, large-scale electrophysiology, and functional ultrasound imaging to determine how these pathways contribute to visually evoked defensive behaviors. Inhibiting either narrow-field or wide-field neurons impaired escape to overhead visual threats, demonstrating that both pathways contribute to escape. Recordings from downstream targets revealed a compact representation in the deeper layers of the superior colliculus that selectively distinguished visual stimuli associated with escape, whereas the lateral posterior thalamus contained a broader representation of visual stimuli. The representation of visual threat in the deeper layers of the superior colliculus predicted escape initiation and vigor in freely moving animals. Across the brain, functional ultrasound imaging showed that visual stimuli recruited distinct patterns of activity that could not be explained by response magnitude alone or by the contribution of single cell-types. Together, these results reveal a cooperative circuit architecture in which parallel pathways carry distinct but complementary signals that jointly support visually evoked escape.

## Introduction

Rapid defensive responses require the brain to transform sensory evidence of threat into appropriate actions. In mammals, the superior colliculus (‘colliculus’) is central to this transformation. It extracts behaviorally relevant features from visual inputs and distributes this information across parallel pathways associated with distinct defensive behaviors (Basso and May, 2017; Cang et al., 2024; Hoy and Farrow, 2025). This organization raises fundamental questions about how visual information is distributed across collicular pathways and whether and how these pathways interact to generate different defensive behaviors.

The colliculus is a layered subcortical structure whose superficial layers receive direct retinal input and contain genetically and morphologically distinct cell types that route visual signals to different downstream targets (Calzoni et al., 2026; Gale and Murphy, 2018, 2014; Li et al., 2023; Reinhard et al., 2019; Sans-Dublanc et al., 2021; Shang et al., 2018; Whyland et al., 2020). Two prominent output streams arise from narrow-field and wide-field neurons. Narrow-field neurons project prominently to the deeper layers of the superior colliculus, a premotor region implicated in orienting and defensive responses, whereas wide-field neurons project to the lateral posterior nucleus of the thalamus, which links collicular visual signals to the amygdala, striatum and broader thalamocortical networks (Calzoni et al., 2026; Li et al., 2023; Sans-Dublanc et al., 2021; Shang et al., 2018; Wei et al., 2015; Wu et al., 2023). Optogenetic activation experiments suggest that these pathways are associated with distinct defensive behaviors, with narrow-field-related circuits promoting escape, and wide-field-related circuits promoting stopping or freezing (Sans-Dublanc et al., 2021; Shang et al., 2018; Wei et al., 2015; Wu et al., 2023).

These findings support a view in which collicular output pathways act as dedicated command lines for specific defensive actions. However, several observations argue that this account is incomplete. Narrow-field and wide-field neurons both respond to a wide variety of behaviorally relevant visual events, including looming and sweeping stimuli, which can trigger escape or freezing (De Franceschi et al., 2016; Kühn et al., 2025; Lee et al., 2020; Li et al., 2023; Relota et al., 2025; Wu et al., 2023; Yilmaz and Meister, 2013). Activation of different collicular cell-types, including wide-field neurons, have been shown to activate vast overlapping networks across the entire brain (Sans-Dublanc et al., 2021). In addition, natural defensive actions are not determined by stimulus identity alone, but depend on behavioral state and context (Daviu et al., 2020; Fratzl et al., 2021; Li et al., 2023; Ritter et al., 2024). These observations suggest that narrow-field and wide-field pathways may not independently specify defensive actions but instead carry complementary information that is combined to generate appropriate behavioral responses. Whether these pathways act independently or cooperatively during visually evoked escapes remains unclear.

Here we address this question by combining cell-typespecific chemogenetic inhibition in freely moving mice with large-scale electrophysiology, optogenetics and functional ultrasound imaging (fUSI). We find that inhibiting either narrow-field projections, which project to the dSC, or wide-field projections, which project to the LP, impairs escape to overhead visual threats, and that the encoding of visual information in the dSC and LP use distinct strategies. Narrow-field neurons contribute to a compact, low-dimensional representation of threat in the deeper layers of the superior colliculus, whereas wide-field neurons support a broad high-dimensional representation of visual stimuli in the lateral posterior thalamus. Brain-wide imaging further reveals stimulus-specific patterns of recruitment, and threat-selective activity in the deeper layers of the superior colliculus predicts escape initiation and vigor. Pathway-specific perturbations differentially alter these representations. Together, these results reveal a cooperative circuit architecture in which parallel collicular output pathways carry distinct but complementary signals that jointly support visually evoked escape.

## Results

### Narrow- and wide-field neurons are each necessary for visually evoked escapes

We asked whether narrow-field and wide-field neurons make distinct contributions to visually evoked escape and whether these contributions depend on stimulus identity. Narrow-field or wide-field neurons were targeted by injecting a Cre-dependent AAV expressing the inhibitory designer receptor exclusively activated by designer drugs (DREADD) hM4Di into the superficial superior colliculus of Grp-Cre or Ntsr1-Cre mice, respectively (Fig. 1A, B). On test days, mice received CNO or saline before entering an open-field arena containing a shelter beneath an overhead display (Fig. 1C). We presented black looming stimuli, which reliably evoke defensive responses, together with three control stimuli matched for contrast or changes in stimulus size (Daviu et al., 2020; De Franceschi et al., 2016; Evans et al., 2018; Li et al., 2023; Yilmaz and Meister, 2013). Under control conditions, black looming evoked escape in 74% ± 6% of trials (mean ± s.e.m.; N = 29 mice).

**Figure 1.**
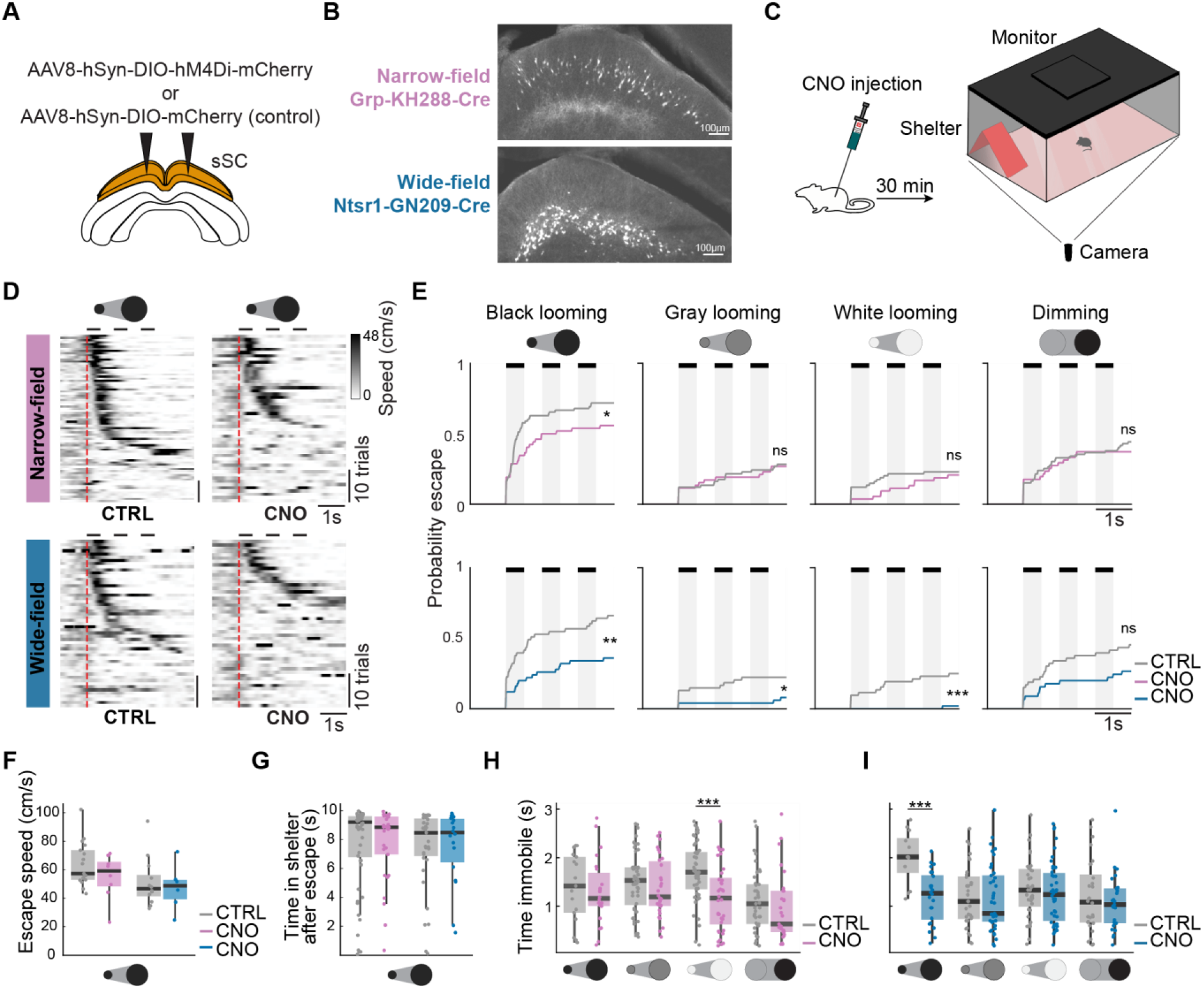
Inhibition of narrow-field and wide-field neurons leads to impairment in escaping behavior. **A**. Schematic of injection strategy. **B**. Fluorescence expression pattern of the AAV carrying hM4Di in Cre mouse lines labeling narrow-field and wide-field neurons. **C**. Schematic of open-field setup, including red transparent floor, an overhead visual display, camera for behavioral tracking, and shelter. **D**. Heatmaps of the speed of the mice for all trials (N = 51 NF CNO, 45 WF CNO, 74 NF CTRL, 50 WF CTRL trials) across all mice (N = 11 NF CNO, 11 WF CNO, 17 NF CTRL, 12 WF CTRL mice) in response to a black looming stimulus. Trials are sorted by time of escape onset. Red dashed line indicates stimulus onset, and three black lines at the top indicate the three presentations of the looming stimulus. **E**. Cumulative probability of escapes. Top: Narrow-field inhibition vs controls. Bottom: Wide-field inhibition vs controls. *p<0.05 **p<0.01, ***p<0.001, log-rank test. **F**. Peak speeds of escapes. p > .05, Mann-Whitney U test. **G**. Time spent in the shelter after an escape bout, maximum time measured is 10 s. p > .05, Mann-Whitney U test. **H**. Immobility time. Narrow-field inhibition vs controls. ***p < 0.001, Mann-Whitney U. test. **I**. Immobility time. Wide-field inhibition vs controls. ***p < 0.001, Mann-Whitney U. test.

Inhibition of either narrow-field or wide-field neurons reduced escape probability in a stimulus dependent manner. Inhibiting narrow-field neurons reduced escape to black looming without affecting escape responses to the other stimuli (Fig. 1D, E; Fig. S1A). In contrast, inhibiting wide-field neurons reduced escape probability across a broader range of visual stimuli (Fig. 1E; Fig. S1B). Neither manipulation altered peak running speed or the time spent in the shelter following escape (Fig. 1F, G), indicating that the ability to execute an escape remained intact and suggesting that narrow-field and wide-field neurons influence the initiation of escape rather than its motor execution.

While inhibition of narrow-field and wide-field neurons reduced the probability of escape, this did not reflect a general loss of responsiveness to visual stimuli. We quantified stimulus-evoked immobility to determine whether the reduction in escape reflected a general decrease in behavioral responsiveness. Narrow-field inhibition altered immobility only in response to white looming, whereas wide-field inhibition altered immobility only in response to black looming; neither manipulation broadly suppressed immobility across the stimulus set (Fig. 1H, I). Together, these results show that narrow-field and wide-field neurons both contribute to escape initiation but do so with different stimulus dependence.

### Targets of narrow- and wide-field neurons emphasize distinct visual features

Narrow- and wide-field neurons both contribute to visually evoked escape, but their contributions differ with stimulus identity. Consistent with distinct functions, selective activation of each population also elicits different behavioral responses (Gale and Murphy, 2014; Hoy et al., 2019; Sans-Dublanc et al., 2021; Wu et al., 2023). We therefore asked whether these pathways convey redundant visual signals or route distinct visual features. Using Neuropixels probes, we performed high-density extracellular recordings in awake, head-fixed mice. We recorded the visual response properties of neurons in the superficial superior colliculus (sSC; 244 cells from 24 mice) and in the principal downstream targets of narrow-field and wide-field cells: the deeper layers of the superior colliculus (dSC; 479 cells from 24 mice) and lateral posterior nucleus of the thalamus (LP; 440 cells from 12 mice), respectively (Fig. 2A–C). Recording sites and regional boundaries were verified using histology, current-source density analysis, and visual response properties (Fig. S2).

**Figure 2.**
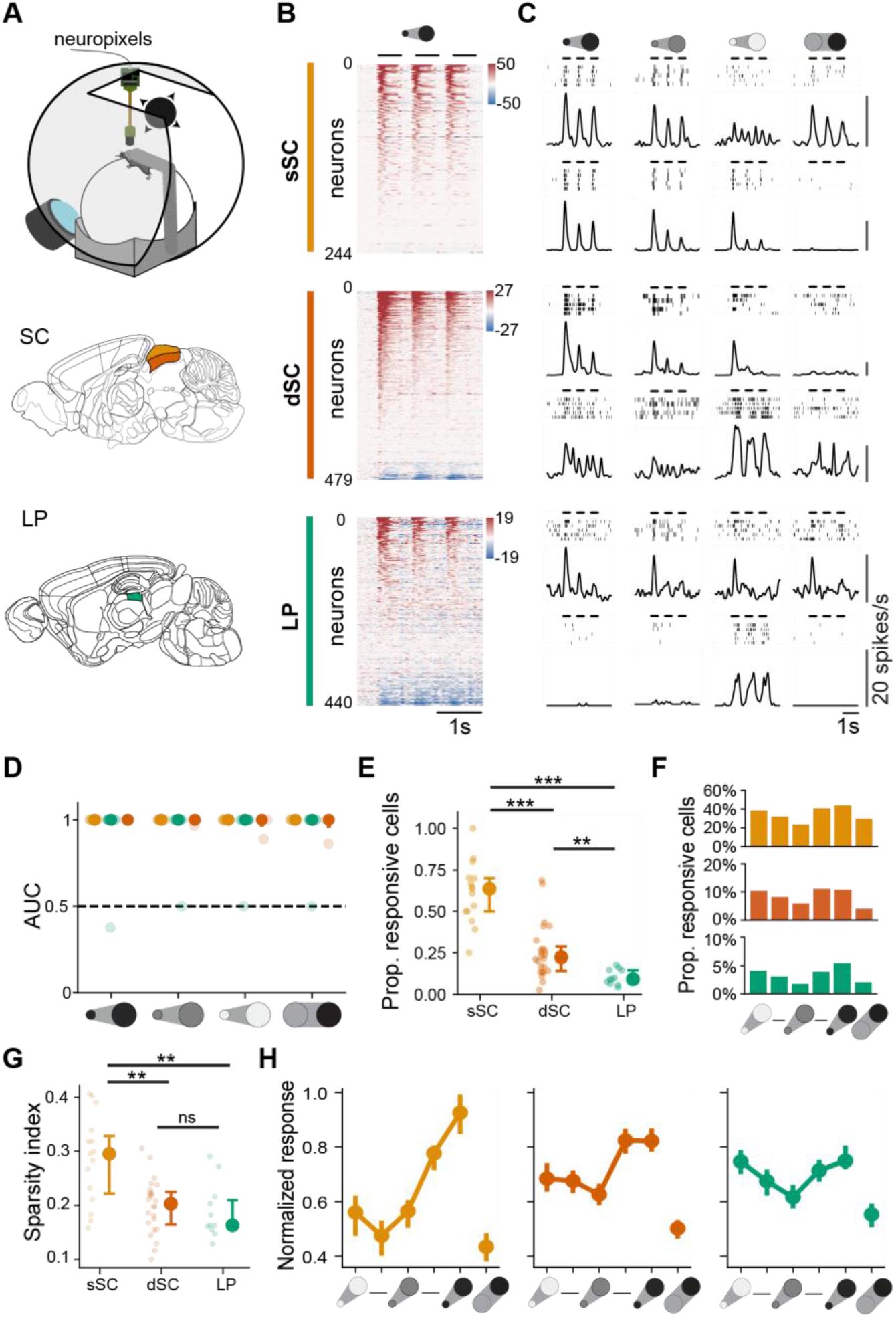
dSC and LP populations detect all stimuli but have different stimulus preferences. **A**. Setup for head-fixed Neuropixels recordings in the superior colliculus (SC) and lateral posterior nucleus (LP) during visual stimulation. **B**. Heatmap of baseline-subtracted neural activity of visually responsive neurons to three repetitions of black looming, ordered by response magnitude. Color bar indicates firing rates (spikes/s). **C**. Example responses of single units to visual stimuli. Left to right: 100% black looming, 25% black looming, 100% white looming, 100% black dimming. **D**. Detection of each visual stimulus decoded by a linear support vector machine (SVM). Dashed line: chance (AUC = 0.5). Transparent markers: individual mice (N = 24 sSC, 24 dSC, 12 LP). Solid marker: median ± IQR. **E**. Proportion of visually responsive neurons across regions (zeta test; p<0.05). Median ± IQR across mice. p_sSC-dSC_ = 1.58 × 10^−5^; p_sSC-LP_ = 2.08 × 10^−5^; p_dSC-LP_ = 1.30 × 10^−3^, Twosided Mann-Whitney-U test. **F**. Proportion of responsive cells across stimuli. Looming contrasts: 100% white, 60% white, 25% black, 60% black, and 100% black. **G**. Adjusted Treves-Rolls lifetime sparsity index averaged per recording. Median ± IQR across mice. p_sSC-dSC_ = 1.91 × 10^−3^; p_sSC-LP_ = 3.09 × 10^−3^; p_dSC-LP_ = 0.42, Mann-Whitney-U test. **H**. Normalized stimulus preferences (maximum response across stimuli) for all visually responsive neurons. Mean ± s.e.m.

Visual stimuli remained detectable from the activity of neural populations, even as visually responsive neurons became progressively less common in downstream regions. To quantify how reliably population activity signaled the presence of a visual stimulus, we trained a support vector machine to distinguish stimulus-evoked activity (Fig. 2D). Looming and dimming stimuli could be reliably detected from neural activity in all three regions, despite marked differences in the fraction of visually responsive neurons. Visual responsiveness decreased from 64% of neurons in the sSC to 22% in the dSC and 9% in LP (Fig. 2E). Low-contrast looming and dimming recruited the fewest neurons in each region (Fig. 2F). Among visually responsive neurons, responses also became less selective across the stimulus set in downstream regions. The adjusted Treves-Rolls sparsity index decreased from the sSC to the dSC and LP (SI = 0.29, 0.21, and 0.17, respectively; Fig. 2G).

Despite this shared capacity to detect stimulus presence, the neural activity in the dSC and LP emphasized different visual features. dSC neurons responded more strongly to black than to white looming, whereas LP neurons responded more strongly to high-than to low-contrast looming (Fig. 2H). Dimming produced the weakest responses across all three regions. These response profiles indicate distinct transformations of visual information in the two downstream regions. dSC preferentially represented the polarity of looming stimuli, including the black looming stimulus that most reliably evoked escape, whereas LP responses were more strongly modulated by stimulus contrast.

### Population geometry of responses to visual threat differs between the dSC and LP

Visual responses occupied a more compact lower-dimensional neural activity space in the dSC than in LP (Fig 3). We quantified the effective dimensionality of stimulus-evoked neural activity using the participation ratio (PR). dSC responses occupied less than half as many dimensions as LP responses (PRdSC = 5.87 [95% CI: 4.89, 7.14]; PRLP = 13.88 [95% CI: 11.17, 16.73]; p < 0.001, two-tailed permutation test; Fig. 3E; Fig. S3C).

**Figure 3.**
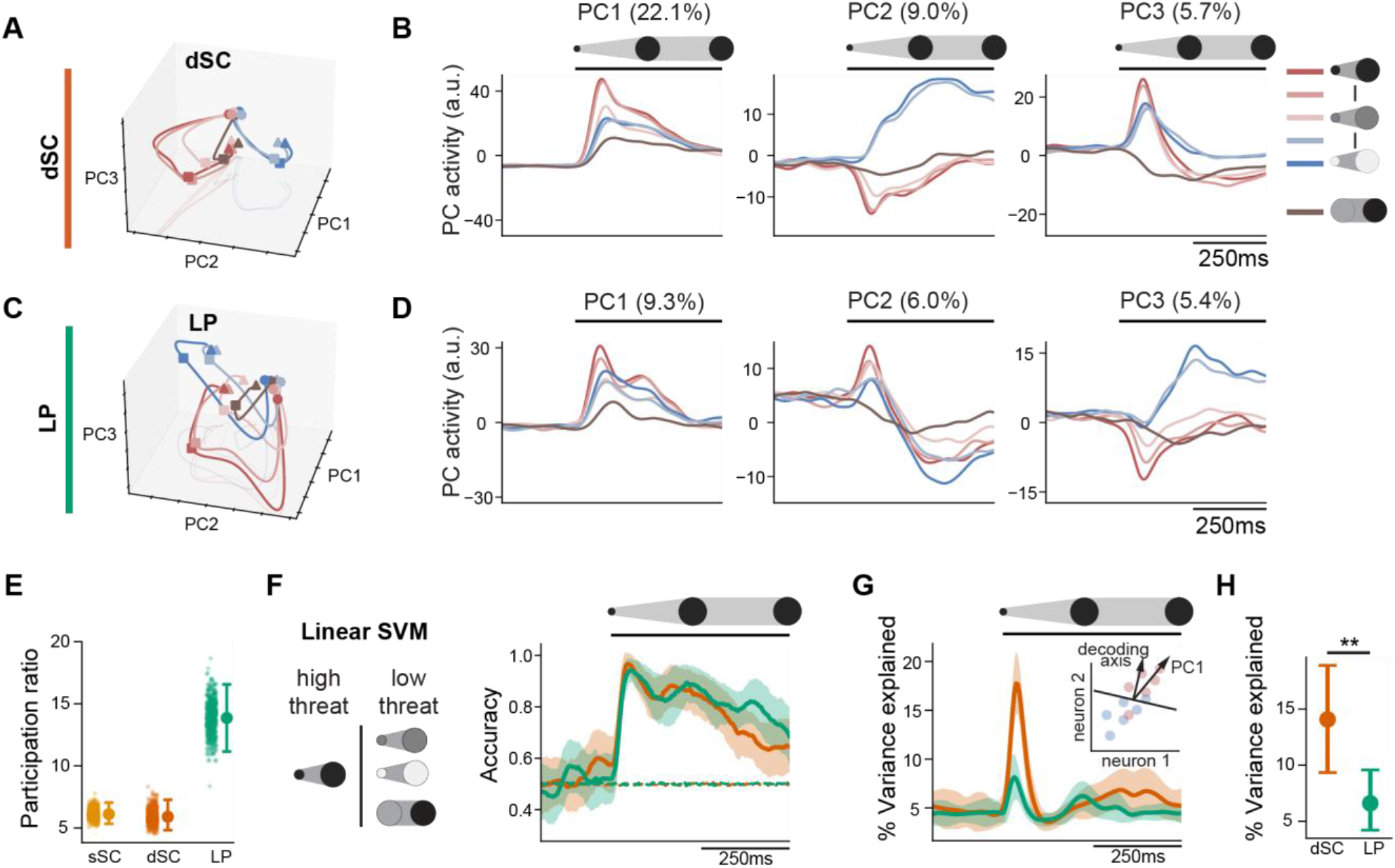
The dSC and LP populations encode stimuli differently. **A,C**. First three principal components based on trial-averaged activity of all cells in all dSC (A) and LP (C) recordings. **B,D**. First two principal components across time for the looming stimuli across contrasts and dimming stimulus in the dSC (B) and LP (D). Looming contrasts: 100% black, 80% black, 40% black (gray), 50% white, and 100% white. **E**. Cross-validated participation ratio (PR) as an estimate of effective dimensionality. Point-and-whisker: median ± 95% confidence interval of bootstrapped distribution (PR_sSC_ = 6.10 [95% CI: 5.30, 7.14], PR_LP_ = 13.88 [95% CI: 11.17, 16.73], PR_dSC_ = 5.87 [95% CI: 4.89, 7.14]) **F**. Left: Linear support vector machine trained to discriminate black looming versus gray looming, white looming and dimming. Right: Decoding accuracy over time. Mean ± s.e.m.. Dashed lines are mean decoding accuracy with shuffled trials. **G**. Percentage of variance in data explained by the decoding axis in dSC and LP. Mean ± s.e.m.. Inset is an illustration of the decoding axis alignment with data variance. **H**. Mean percent signal variance explained in 25 ms – 75 ms post stimulus time window. Point-and-whisker: mean ± 95% confidence interval of bootstrapped distribution. Permutation tests were used to compare the variance explained in each region against a null distribution (p < 0.01 for both dSC and LP). Regions were compared using the bootstrapped distribution of their difference (difference = 7.72% [95% CI: 2.72%, 13.35%]; p = 0.004).

Despite these different geometries, both regions contained information that distinguished the visual stimulus most strongly associated with escape from stimuli that rarely evoked escape. Based on escape probability (Fig. 1), we classified high-contrast black looming as the high-threat stimulus and low-contrast black looming, white looming, and dimming as low-threat stimuli. Linear support-vector machines trained on pseudo-populations reliably discriminated these behaviorally defined stimulus categories from activity in both the dSC and LP, with decoding performance peaking approximately 30 ms after stimulus onset (Fig. 3F; Fig. S3D).

Information distinguishing the high-threat stimulus was concentrated along a dominant population dimension in the dSC but distributed across dimensions in the LP. We measured the fraction of population variance captured by the support vector machine (SVM) threat-decoding axis to determine where threat information lay within each population geometry. The threat axis captured a larger fraction of total variance in dSC than in LP (Fig. 3G and H). In dSC, the threat-decoding axis aligned closely with the first principal component, indicating that threat-related activity was concentrated along a dominant dimension of population variance. In LP, the threat-decoding axis projected onto multiple lower-variance principal components (Fig. S3E).

### Narrow- and wide-field neurons differentially contribute to downstream targets

To determine how narrow- and wide-field cells contribute to the representation of visual information in the dSC and LP, we optogenetically identified neurons receiving input from each collicular population. We expressed ChR2 in either narrow-field or wide-field neurons using GRP×ChR2 and NTSR1xChR2 mice and recorded neural activity during optogenetic stimulation (Fig 4A, S6A). Putative targets of each cell type were identified based on their response latency and reliability. We defined secondary cells as those that responded within 9 ms of the stimulation, at least 10% of time (Fig 4B). These neurons occupied a narrow band immediately below the optic layer, corresponding to the deeper layers of the superior colliculus (Fig 4C).

**Figure 4.**
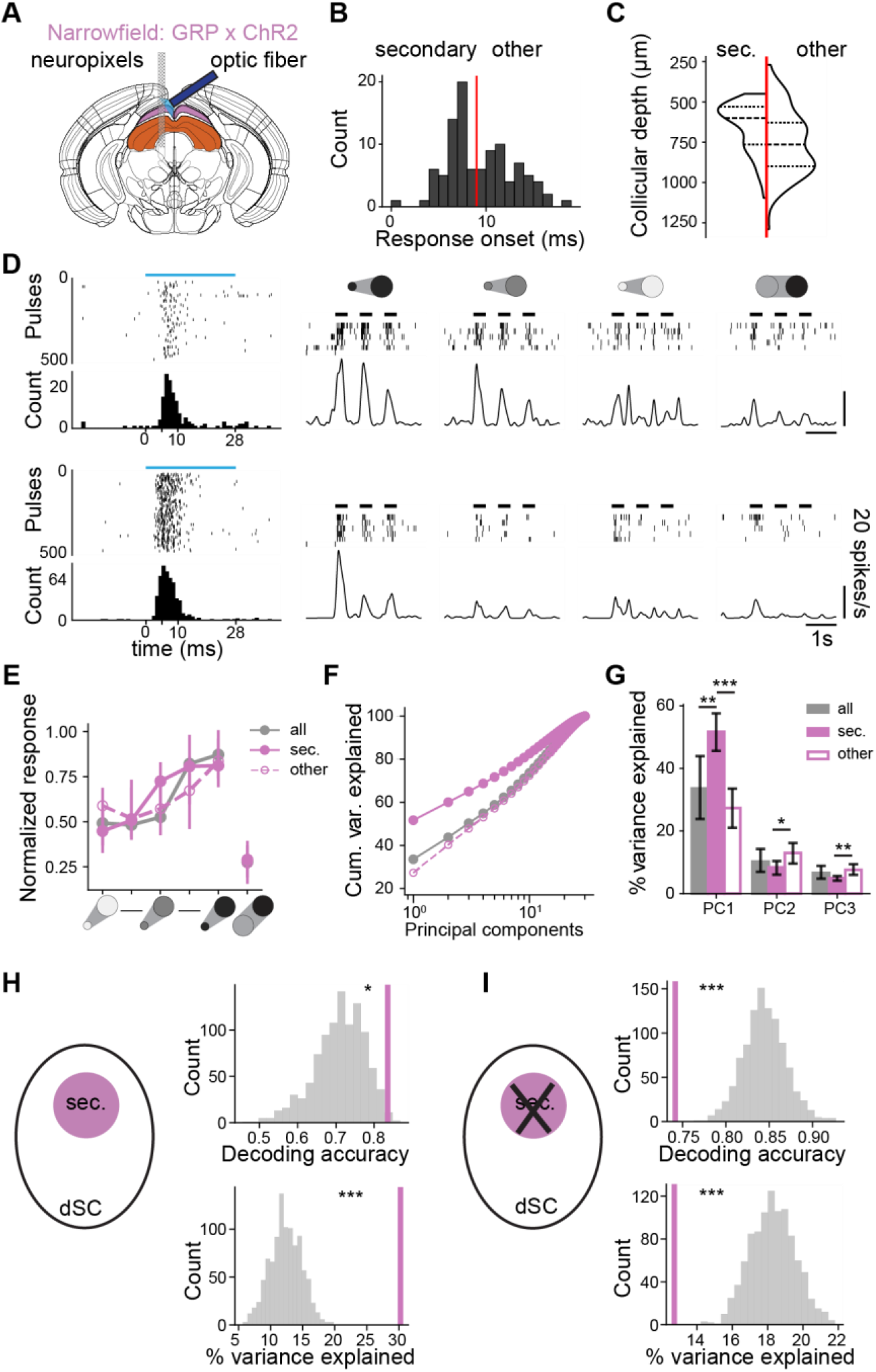
Narrow-field cells drive threat representations in the dSC. **A**. Schematic of the experimental setup. **B**. Distribution of response onset latencies to optogenetic stimulation. Red line at 9 ms. **C**. Anatomical distribution of putative secondary and other optotagged cells across the depth of the colliculus, from the surface of the sSC. **D**. Two representative secondary cells. Left: responses to 500 optogenetic stimulation pulses (28 ms duration, blue bar). Right: raster plots and corresponding peristimulus time histograms. **E**. Stimulus preferences for visually responsive dSC cells, normalized to the maximum response (mean ± s.e.m). **F**. Cumulative variance explained across principal components for the complete population of dSC neurons, secondary optotagged, and non-secondary optotagged neurons, respectively. **G**. Variance explained by the first three PCs, mean ± 95% bootstrapped confidence interval (CI). PC1: p_all-sec_ = 0.002, p_sec-other_ < 0.001; PC2: p_all-sec_ = 0.024; PC3: p_secother_ = 0.001. **H**. Comparison of the decoding accuracy (top) and variance explained by the decoding axis (bottom) of secondary optotagged cells (purple line) versus sizematched subsets of other dSC neurons (p_decoding_ = 0.011, p_variance_ < 0.001; N = 6 mice, N_sec_ = 42 cells, N_other_ = 50, N_all_ = 354). **I**. Comparison of the decoding accuracy (top) and variance explained by the decoding axis (bottom) when excluding secondary cells (purple line) versus excluding size-matched subsets of other dSC neurons (p_decoding_ < 0.001, p_variance_ < 0.001; N = 6 mice; N_sec_ = 42 cells, N_other_ = 50, N_all_ = 354).

Neurons in the dSC receiving direct input from narrow-field neurons, i.e. putative secondary neurons, were enriched for the dominant visual responses in the dSC. Visually responsive secondary cells showed the same preference for black looming as late-responding optogenetically activated neurons and the broader dSC population (Fig. 4E). However, visual responses were more common among secondary than late-responding cells (40.6% versus 20.3%; Fig. S5). Population activity also differed in its organization. The first principal component accounted for more variance among secondary cells than in the broader dSC population (Fig. 4F and G). Because this component captured the dominant visually evoked response in dSC (Fig. 3B), narrow-field-recipient neurons were preferentially aligned with the major visual dimension of the dSC population.

Additionally, neurons receiving narrow-field input contain a disproportionate fraction of the sensory threat information represented in dSC. We tested whether secondary cells contained sufficient information to distinguish high- from lower-threat stimuli and what fraction threat-related information they accounted for in the population. A decoder trained exclusively on secondary cells reliably classified threat and outperformed equivalently sized subsets of non-optotagged dSC neurons (Fig. 4H, top). The threat-decoding axis captured more population variance among secondary cells than the rest of the neural population (Fig. 4H, bottom). Conversely, computationally removing secondary cells from the general dSC population reduced threat-decoding performance and decreased the variance captured by the threat axis (Fig. 4I).

On the other hand, while wide-field-recipient neurons broadly reflected the visual representation within the LP population, they did not form a population enriched for a particular visual dimension. We identified putative wide-field-recipient LP neurons using the same latency and response-probability criteria (Fig. S6B). Their visual stimulus preferences were similar to those of the remaining LP population (Fig. S6D), and the distribution of the variance of the neural population across principal components did not differ between recipient and non-recipient neurons (Fig. S6E, F). Together, these results indicate that narrow- and wide-field inputs contribute differently to their downstream representations: narrow-field input preferentially targets dSC neurons enriched for sensory threat information, whereas wide-field-recipient neurons reflect the broader visual representation distributed across LP.

### Representation of threat in the dSC predicts escape initiation and vigor

The preceding analyses demonstrated that neural activity in the dSC represents the sensory threat value of visual stimuli under head fixation. We asked whether this representation predicts escape behavior in freely moving animals. First, we tested whether neural activity in the dSC and LP activity predicts escape. Second, we tested whether the encoding of sensory threat generalizes across conditions, head-fixed versus freely moving. Using chronically implanted Neuropixels 2 probes, we recorded neural activity from the same mice during freely moving and head-fixed sessions (Steinmetz et al., 2021) (Fig. 5A, B). During freely moving sessions, mice viewed overhead visual stimuli in an arena containing a shelter. We subsequently presented the same stimuli under head fixation to characterize the visual response properties of the recorded neurons. Joint spike sorting across recording conditions allowed us to track neurons between freely moving and head-fixed sessions. Implanted mice reliably escaped from black looming stimuli (Fig. 5C), and recorded neurons responded robustly in both settings (Fig. 5D).

**Figure 5.**
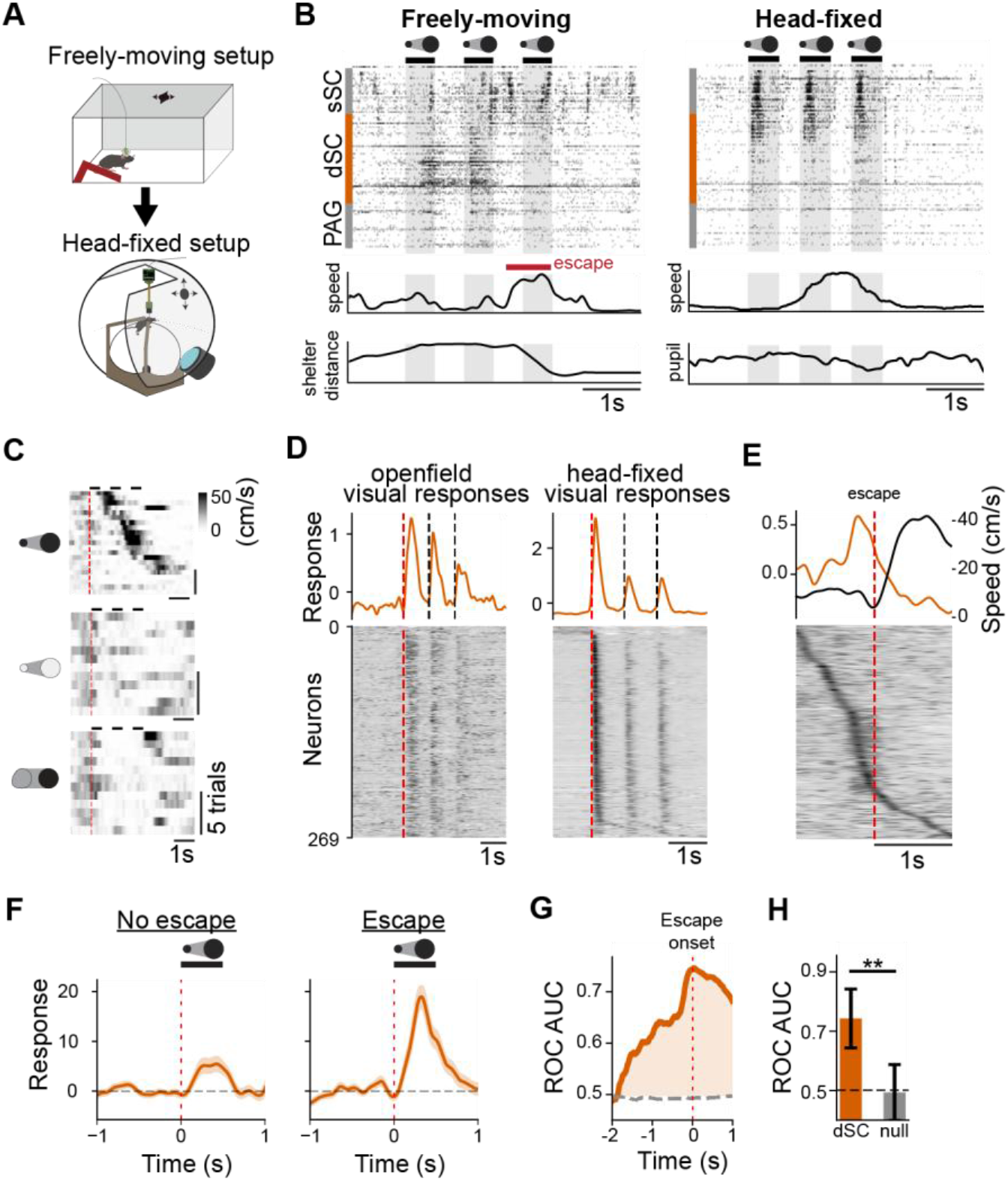
dSC activity precedes and predicts escape responses. **A**. Schematic of the two-stage recording paradigm. Animals were first exposed to visual stimuli in an open-field setup and subsequently presented the same visual stimuli during headfixed recordings. **B**. Representative spike rasters of recorded cells across collicular depth in both the freely moving and headfixed setups. Bottom traces show simultaneously recorded running speed, distance to shelter, and pupil diameter. Red horizontal bar indicates an escape response. **C**. Running speed heatmaps across trials aligned to stimulus onset (red dashed line) for black looming, white looming, and dimming stimuli. **D**. Mean z-scored population activity (top) and z-scored response heatmaps (bottom) aligned to visual stimulus onset in open-field (left) and head-fixed (right) setups. Averaged across stimuli. Cells are sorted by peak response latency in the head-fixed setup (269 cells, N = 4 mice). **E**. Mean z-scored population activity (top) and z-scored response heatmaps (bottom) aligned to escape onset in the open-field setup. Cells are sorted by peak response latency. **F**. Mean dSC population response to black looming stimuli on escape trials versus non-escape trials. Zoomed in on first loom repetition indicated by overhead bar. Shaded areas represent mean ± SEM across cells. p = 5.54 × 10-10, Mann – Whitney U Test; N = 21 trials). **G**. Receiver operating characteristic area under the curve (ROC-AUC) over time aligned to escape onset. Gray dashed line denotes chance-level AUC (shuffled null distribution). **H**. Mean ROC-AUC values during the 100 ms preceding escape onset. Mean ± SD of bootstrapped trials. Permutation test was used to compare the dSC performance against a null distribution (p = 0.004; N = 34 trials).

We found that dSC activity preceding movement predicted whether mice escaped. Black looming evoked stronger dSC responses on escape compared to non-escape trials (Fig. 5F). Activity increased during stimulus expansion, reached its maximum around escape onset, and declined after movement began (Fig. 5E). Individual neurons showed heterogeneous temporal profiles, including a prominent population whose activity peaked immediately before movement onset (Fig. 5E). Consistent with these single-neuron dynamics, ROC discrimination between escape and non-escape trials increased progressively during stimulus expansion and peaked around escape onset (Fig. 5G and H). These dynamics are consistent with previous calcium imaging measurements of dSC activity during visually evoked escape (Evans et al., 2018). Exploratory recordings in LP (N = 2 mice; four sessions) revealed visual responses during freely moving behavior but did not show the same relationship with escape initiation (Fig. S4).

The representation of sensory threat in the population activity was conserved across animals. To determine whether the head-fixed representation of sensory threat could be transferred to freely moving behavior, we first aligned head-fixed population activity across animals using multiset canonical correlation analysis (multiCCA) (Kettenring, 1971; Vinod, 1976). This analysis defined a shared latent space into which we subsequently projected freely moving activity from the same neurons (Fig. 6A; Methods). Based on the effective dimensionality of dSC activity, we retained five latent dimensions. The first four dimensions aligned significantly across animals, whereas the fifth did not (CV1–CV4, p < 0.05; CV5, p = 0.27; stimulus-shuffled permutation test; Fig. 6B). We then trained a linear support-vector machine in this shared space to distinguish high-threat black looming from lower-threat white and gray looming and dimming stimuli. In leave-one-session-out cross-validation, a classifier trained on N−1 sessions accurately classified threat category in the held-out session from the mean response 25–75 ms after stimulus onset (accuracy = 92.5%; Fig. 6C, D).

**Figure 6.**
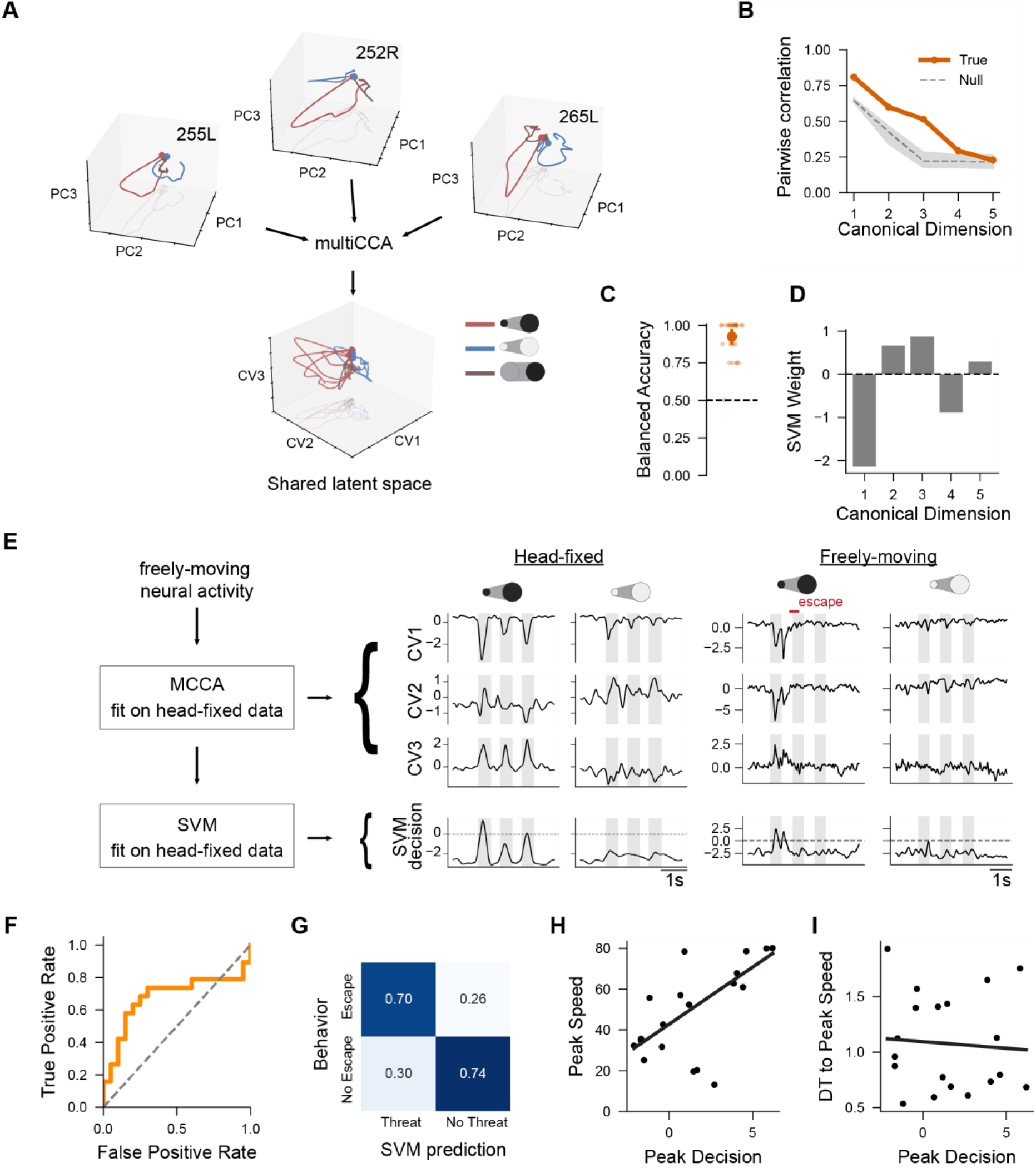
Representation of threat in dSC reliably precedes defensive behaviors. **A**. Schematic of the analytic approach used to align the neural data across animals. (N = 27 mice; 30 sessions) **B**. Assessment of cross-animal alignment via pairwise correlations across canonical variates (CVs). Correlations for the first four dimensions are significantly higher than a null distribution generated by shuffling stimuli (gray dashed line with shaded confidence interval) (CVs 1–4, p < 0.05; CV 5, p = 0.27). **C**. Balanced decoding accuracy of the Support Vector Machine (SVM) classifier across animals using leave-one-out cross validation. The dashed line indicates chance level. Significance was determined using the Wilcoxon test (accuracy = 0.925 [95% CI: 0.871, 0.958]; p=9.49 × 10^−7^). **D**. SVM weights assigned to the canonical variates, illustrating each dimension’s contribution to the decision boundary. **E**. Application of the head-fixed decoder to freely-moving behavior. Left: the multiCCA and SVM model were fit on head-fixed data before being applied to freely-moving neural activity. Right: example trials showing the first three CVs and the SVM decision value during head-fixed and freely-moving settings. **F**. Receiver Operating Characteristic (ROC) curve evaluating the SVM decision boundary’s ability to predict escape responses across classification thresholds (ROC-AUC = 0.679). **G**. Confusion matrix comparing the SVM prediction of threat to the behavioral response (escape vs no escape) using the optimal threshold derived from the ROC curve. **H**. Correlation between the peak SVM decision value prior to an escape bout (calculated within a 3 s post stimulus onset window) and escape magnitude, quantified as the peak running speed. (Pearson’s r = 0.611, p = 0.005). **I**. Correlation between the peak SVM decision value prior to an escape bout and the time to the peak escape speed (Pearson’s r = −0.094, p = 0.731).

The representation of sensory threat defined under head fixation predicted freely moving escape. We projected freely moving activity into the shared latent space and applied the classifier trained exclusively on head-fixed recordings. Classifier decision values were higher before escape (Fig. 6E), and ROC analysis showed that these values reliably predicted escapes (Fig. 6F, G). Pre-escape decision values did not correlate with escape latency (Fig. 6I), but increased with escape speed (Fig. 6H), consistent with calcium imaging experiments (Evans et al., 2018). Thus, representation of visual threat in the dSC identified under head fixation predicts both the initiation and vigor of escape in freely moving animals.

### Visual stimuli recruit distinct, overlapping brain-wide networks

The distinct representations of visual information in the dSC and LP suggest that parallel collicular pathways are embedded within broader networks that process visual threat. We asked how the same visual stimuli recruit activity across the brain, and whether cell-types specific pathways provides a major route through which visual signals engage these distributed networks. Using functional ultrasound imaging (fUSI), we measured brain-wide responses to looming, dimming, and sweeping visual stimuli (Fig. 7A). The three stimuli recruited overlapping but distinct brain-wide sets of brain regions (Fig. 7A–C, E). Across all stimuli, 125 regions were significantly activated, but only 26 responded to all three (Fig. 7A, C). Shared responses included major visual structures, including the superior colliculus (SCs), dorsal lateral geniculate nucleus (LGd), primary and higher visual cortex (VISp, VISl, VISli, VISpl, VISpor), and temporal association cortex (TEa). Common recruitment also extended to hippocampal and regions involved in navigation (CA3d, DGd, ENTm, PAR, POST, ProS, SUB and RSP), as well as structures associated with visually guided innate behaviors, including the lateral posterior thalamus (LP), deeper layers of the superior colliculus (SCd/SCi), periaqueductal gray (PAG), and pretectal nuclei (NOT, PPT) (Fig. 7A, D, E).

**Figure 7.**
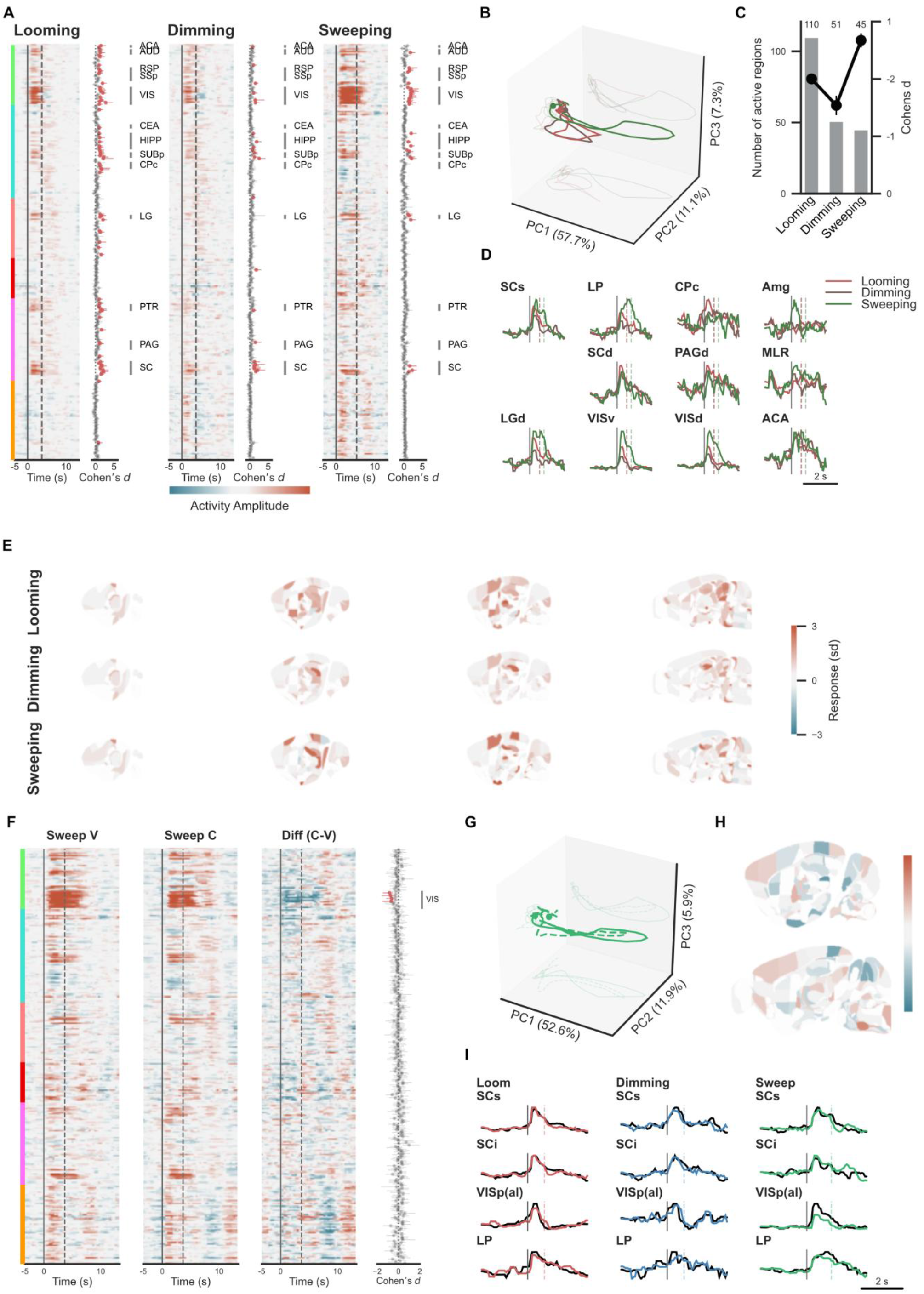
Brain-wide fUSI reveals stimulus-specific visual responses. **A**. Brain-wide fUSI responses to looming, dimming, and sweeping stimuli across atlas-defined regions. Right, response significance for each region. **B**. PCA trajectories of brain-wide activity evoked by each stimulus. **C**. Number of significantly activated regions and corresponding effect size (Cohen’s ***d***) for each stimulus. **D**. Mean stimulus-evoked responses in representative brain regions. Vertical dashed lines indicate the offset of looming and dimming (red) and sweeping stimuli (green). **E**. Anatomical maps of responses to looming, dimming, and sweeping stimuli across representative brain sections. **F**. Brain-wide responses to sweeping stimuli under vehicle (V) and CNO (C) administration, and their difference (C−V). Right, Cohen’s ***d*** for the C−V comparison across regions. **G**. PCA trajectories of brain-wide responses to the two sweeping conditions. **H**. Anatomical maps of response differences between sweeping conditions. **I**. Mean responses in SCs, SCi, VISp(al), and LP to looming, dimming, and sweeping stimuli. Vertical lines indicate stimulus timing.

The most prominent difference was the extent of brain-wide recruitment. Looming activated substantially more regions than dimming or sweeping (110, 51 and 43 regions, respectively; Fig. 7A, C), including 29 regions not activated by either of the other stimuli (Fig. 7A, E). These 29 regions included non-image-forming visual regions (e.g. the intergeniculate leaflet (IGL), ventral lateral geniculate nucleus (LGv), anterior and medial pretectal nuclei (APN, MPT), nucleus of the posterior commissure (NPC), and PRC/SCO), as well as components of the habenulo-interpeduncular network, which included the lateral and medial habenula (LH, MH), several of their associated input regions (LSc, TRS, CA1i, FC, ACAv and caudal caudoputamen subdivisions), and downstream midbrain regions including the interpeduncular nucleus, PAG and midbrain reticular nucleus (MRN) (Fig. 7A, E). Beyond its broader recruitment, looming shared a network with sweeping that was less strongly engaged by dimming. Thirteen regions responded to both looming and sweeping but were not significantly modulated by dimming (Fig. 7A, E). These included higher visual areas (VISal, VISam, VISpm, VISrl), subdivisions of primary visual cortex [VISp(al), VISp(am), VISp(pm)], retrosplenial cortex (RSPagl, RSPd), the anterior pretectal region (APr), and lateral dorsal thalamus (LD). Thus, defensive visual stimuli recruited a distributed set of visuospatial regions in addition to the network shared across all three stimuli.We next asked whether wide-field neurons are required to recruit these distributed visual responses, combining chemogenetic inhibition of wide-field neurons with fUSI. Inhibiting wide-field neurons reduced sweeping-evoked responses in visual cortical areas (Fig. 7F-I), but produced no detectable changes in most other stimulus-evoked responses across the brain (Fig. 7F, G). Thus, despite their importance for visually evoked escape, wide-field neurons were not required for most of the distributed visual responses measured here. This dissociation argues against a model in which the wide-field pathway independently drives a stimulus-specific brain-wide network. Instead, it is consistent with a more selective role in modulating how threat-related activity in the dSC is converted into escape.

### Pathway activity is consistent with a non-linear driver-gate architecture

Given that activity in the dSC strongly represented visual threat, whereas wide-field neurons influenced escape without broadly driving brain-wide visual responses, we asked how the two pathways might interact to control behavior. We therefore evaluated three population readout architectures: competition, linear summation, and a multiplicative driver–gate model (Fig. 8A).

**Figure 8.**
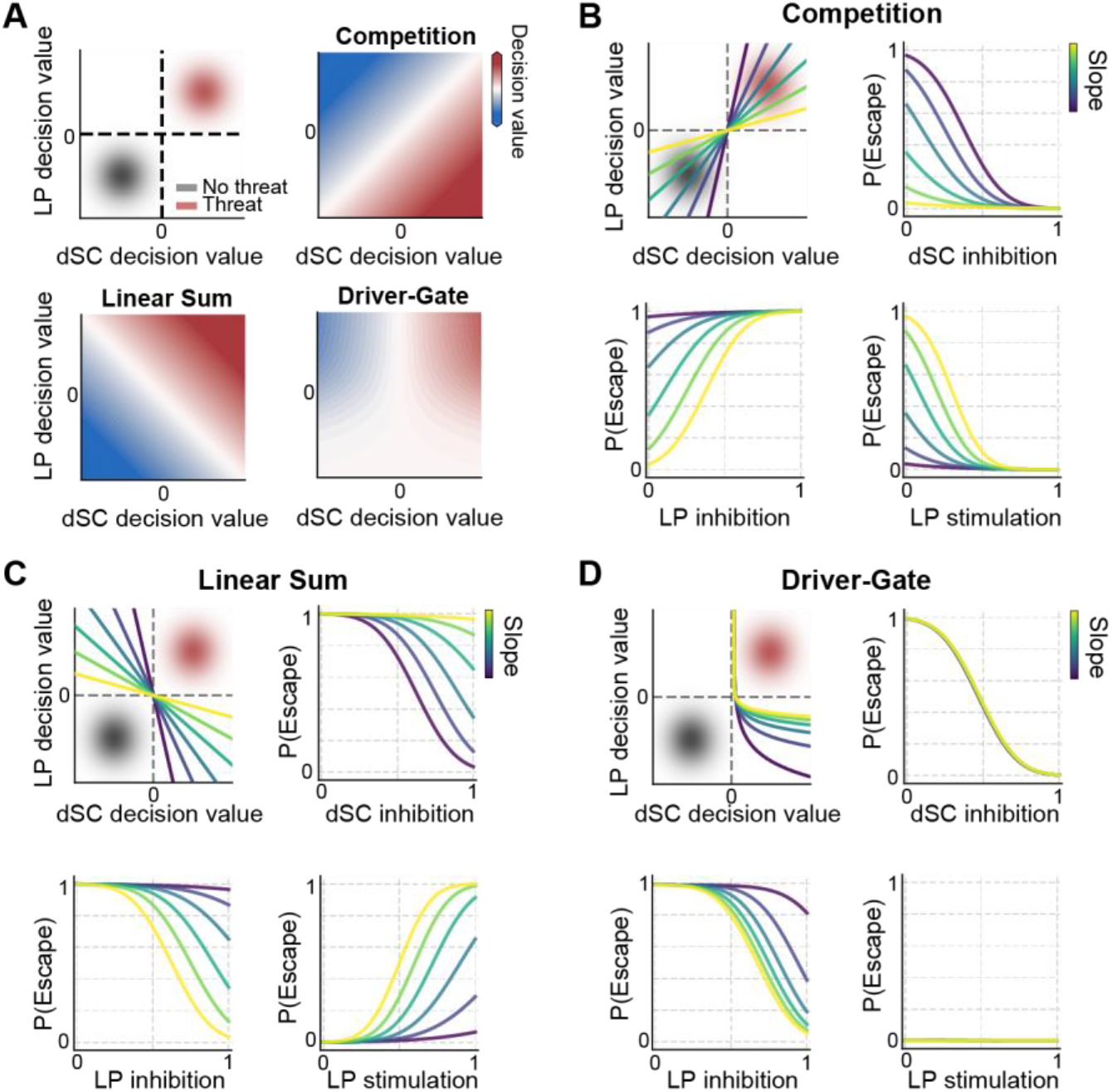
Pathways activity is consistent with a non-linear driver-gate model. **A**. Top left: 2D state space defined by threat decoder decision values of the dSC (x-axis) and LP (y-axis). Decision values for no-threat stimuli (gray) and threat stimuli (red) are modeled as 2D Gaussian probability distributions based on experimental data (Fig. S7A). Top right: Linear Sum model (dSC + LP), Bottom left: Competition model (dSC - LP), Bottom right: Multiplicative Driver-Gate (dSC x σ(LP)). **B-D**. Behavioral simulations evaluating the Competition (B), Linear Sum (C), and Driver-Gate (D) models under simulated neural manipulations. Top left: decision boundaries across a range of LP slopes. Top right: simulated escape probability as a function of increasing dSC inhibition. Bottom left: simulated escape probability as a function of increasing LP inhibition. Bottom right: simulated escape probability as a function of increasing LP stimulation.

In each model, threat information from the two pathways is combined in a two-dimensional space whose axes represent the threat evidence carried by each region’s population activity (quantified as the signed distance from the threat decoder’s decision boundary). Within this space, we simulated Gaussian distributions representing high-threat (black looming) and low-threat (gray looming, white looming, or dimming) stimuli based on our in vivo recordings (Fig. S7A). For each architecture, we tested a set of decision boundaries and compared the predicted behavioral consequences of simulated manipulations with our experimental observations.

Each of the three model architectures was consistent with a driver-like contribution from dSC. Suppressing dSC activity reduced the dSC decision value and decreased predicted escape probability, whereas increasing dSC activity shifted responses toward the escape region of state space (Fig. 8B-D; Fig. S8). These predictions matched the effects of experimental manipulations of the dSC pathway.

In contrast, competitive and linear summation architectures failed to reproduce the effects of perturbing the LP pathway. In the competitive model, suppressing LP removed a subtractive contribution to the downstream readout and therefore increased predicted escape probability (Fig. 8B; Fig. S8A). This prediction was opposite to the reduced escape probability observed after inhibition of wide-field neurons (Fig. 1). In the linear summation model, increasing LP activity shifted the state vector toward the escape boundary and predicted increased escape probability (Fig. 8C; Fig. S8A). This was inconsistent with previous observations that optogenetic activation of the LP pathway induces stopping but does not evoke escape (Sans-Dublanc et al., 2021).

A multiplicative driver-gate architecture reproduced the behavioral effects of perturbing the LP and dSC. In this model, dSC and LP interact nonlinearly to generate an L-shaped decision boundary (Fig. 8D; Fig. S8A). Increasing LP activity alone moves the population state parallel to the escape boundary and therefore does not trigger escape. Conversely, reducing LP activity decreases the probability that a concurrent dSC threat signal crosses the escape threshold. Escape therefore requires both sufficient dSC drive and permissive LP activity. The observed effects of pathway activation and inhibition are consistent with a model in which dSC provides the principal escape drive while LP gates the transformation of that signal into action.

## Discussion

Previous studies have associated narrow-field and wide-field neurons with distinct defensive behaviors, raising the possibility that these cell types form dedicated pathways that independently select behavioral responses. Our results suggest a different circuit organization. Inhibiting either population reduced visually evoked escape, demonstrating that both contribute to the same defensive action. However, the two pathways did not carry the same information. Narrow-field input contributed preferentially to a compact representation of visual threat in the dSC, while wide-field input to the LP contributed to the broader representation of visual stimuli there. Together, our data suggest that parallel output pathways of the colliculus cooperate to support visually evoked escape while making distinct contributions to the transformation of visual information into behavior.

The cooperative contribution of these pathways appears at odds with the distinct defensive behaviors produced by their selective activation. Activating narrow-field- and wide-field-related circuits can evoke different responses, including escape, stopping or freezing (Sans-Dublanc et al., 2021; Shang et al., 2018; Wei et al., 2015; Wu et al., 2023). These findings have motivated models in which parallel collicular pathways act as dedicated channels for different defensive actions. Such models predict that inhibiting one pathway should selectively impair its associated behavior or bias behavior in some directed manner. Instead, inhibiting either narrow-field or wide-field neurons reduced escape, demonstrating that the behaviors produced by activating individual pathways do not necessarily define their contributions during visually evoked behavior. Instead, each pathway contributes to escape despite producing different behavioral effects when activated alone. Together, these findings suggest that parallel, cell-type-specific collicular pathways do not independently specify defensive actions but instead cooperate through functionally distinct contributions to visually evoked behavior.

Narrow-field input links the representation of visual threat in dSC to escape initiation. The dSC has an established role in visually guided defensive behavior, providing input to downstream premotor structures in the PAG involved in escape (Evans et al., 2018; Lee et al., 2020). Here, visual activity in dSC occupied a relatively low-dimensional population space whose dominant dimension distinguished high-threat black looming from dimming or white looming, stimuli that less reliably evoke escape (Yilmaz and Meister, 2013). Narrow-field-recipient neurons were enriched for this representation, indicating that narrow-field input contributes preferentially to the dominant threat-related signal in dSC rather than sampling its activity uniformly. A decoder defined from responses in head-fixed animals predicted whether freely moving animals escaped, and its activity before escape covaried with escape vigor. These observations link the sensory organization of narrow-field input to a dSC population signal that precedes behavior. This result extends previous observations that dSC activity increases before visually evoked escape (Evans et al., 2018) by identifying a threat-related population representation that is conserved across animals and predicts subsequent action.

In contrast to the narrow-field pathway, the wide-field pathway contributes to a broader representation of visual information in LP. Threat-related information in LP was distributed across a higher-dimensional space rather than concentrated along a dominant dimension, and wide-field-recipient neurons reflected this broader representation rather than forming a population enriched for threat-related information. In addition, unlike activity in the dSC, in exploratory recordings from the LP, we did not detect the selective increase preceding escape that was observed in the dSC, suggesting that although both pathways were required for visually evoked escape, their downstream activity differed markedly in its relationship to threat and behavior. Previous work shows that responses in wide-field neurons vary with locomotion and that the tectothalamic pathway supplies several visual cortical areas, raising the possibility that this pathway helps combine visual information with behavioral-state signals and distribute it across the wider network (Beltramo and Scanziani, 2019; Brenner et al., 2023; Relota et al., 2025).

Brain-wide imaging further showed that looming, sweeping, and dimming recruited overlapping but distinct sets of brain regions. Inhibiting wide-field neurons selectively reduced sweeping-evoked responses in visual cortical areas, while leaving most other brain-wide responses unchanged. Wide-field neurons therefore contribute to a specific component of cortical recruitment but do not provide the dominant route through which visual signals reach the wider network.

The asymmetric contributions of the two pathways are consistent with a nonlinear driver-gate architecture. A simple independent-pathway model cannot explain why inhibiting either pathway reduces escape, whereas an opponent model predicts that suppressing a pathway associated with stopping should facilitate rather than impair escape. Linear summation also has difficulty reconciling the necessity of wide-field activity with previous observations that activating this pathway does not trigger escape. A multiplicative interaction provides one solution to these constraints. In this model, the threat-related dSC signal provides the principal drive toward escape, whereas activity associated with the wide-field pathway determines whether that drive reaches the threshold for action. Such an interaction reproduces the effects of pathway inhibition and activation without requiring the two pathways to carry equivalent signals. We therefore interpret the wide-field pathway as providing a permissive contribution to escape rather than an independent escape command. The present experiments do not establish the underlying operation as multiplicative, however, and the driver-gate model should be viewed as a circuit hypothesis that accounts for the observed asymmetry between the pathways.

The anatomical implementation of this cooperative interaction remains unresolved. Our perturbations targeted cell types rather than individual projections, and both populations participate in circuits extending beyond the downstream regions examined here. Narrow-field neurons project to the dSC but also send projections to the parabigeminal nucleus, so the behavioral effect of narrow-field inhibition cannot be assigned specifically to the narrow-field → dSC projection. Similarly, the requirement for wide-field neurons does not establish that LP itself provides the permissive signal proposed by the model. The two pathways could influence a common downstream structure, interact through recurrent or feedback circuits, or affect escape through different nodes of a distributed network. Identifying where pathway activity becomes jointly necessary, and testing whether manipulating the proposed interaction changes the relationship between dSC threat signals and behavior, will be required to distinguish among these possibilities.

These findings revise a simple labeled-line account of collicular control of defensive behavior. Parallel output pathways do not necessarily map independently onto distinct actions. Narrow-field and wide-field neurons are both required for visually evoked escape, yet their downstream targets organize visual information differently: dSC concentrates threat-related information in a population signal closely associated with escape, whereas LP maintains a broader representation of visual stimuli. This division of information provides a circuit architecture in which parallel pathways can make complementary contributions to the same behavior. More generally, cooperation between pathways carrying different representations provides a potential mechanism for preserving rapid responses to visual threat while allowing behavioral output to depend on information distributed across the wider network.

## Acknowledgements

We thank: Gioia De Franceschi for her helpful advice on data analysis; Frederique Ooms for her assistance with histology; Alex Calzoni for viral vector production, and Antonio Ortega and Clement Brunner for performing surgeries. This work was supported by funding from FWO grant G091719N and G018324N (K.F.), NIH grant 1RO1EY032101 (K.F.), KU Leuven Research Council grant C14_22_074 (K.F.), VIB TechWatch (K.F., V.B), and FWO fellowships 1197818N/1197820N (A.S.-D.), and 1205421N (N.K.), 11C5119N/11C5121N (A.C.). This paper was typeset with the bioRxiv word template by @Chrelli: www.github.com/chrelli/bioRxivword-template.

## Author contributions

B.N., A.C., A.S.-D., A.U., and K.F. conceived and designed the study. B.N., A.S.-D., A.C., Z.D., C.L., L.H., R.L., A.B., M.G., and N.K.

performed the experiments. B.N., A.S.-D., A.C., G.M., P.J.G. and

K.F. analyzed the data. A.P. designed experimental setups. B.N., A.S.-D., A.C., and K.F. wrote the manuscript. A.U., K.F., V.B. and

P.J.G. supervised and funded the work.

## Competing interest statement

We have no competing interest.

## Data availability

The datasets generated in the current study are available from the corresponding author upon reasonable request.

## Methods

### EXPERIMENTAL MODEL AND STUDY PARTICIPANT DETAILS

#### Mice

All experimental procedures were approved by the Ethical Committee for Animal Experimentation (ECD) of the KU Leuven and followed the European Communities Guidelines on the Care and Use of Laboratory Animals (004–2014/EEC, 240–2013/EEC, 252–2015/EEC, 165-2018/EEC, 055-2023/EEC, 146-2023/EEC). Male and female adult (2-5 months old) transgenic mice were used in our experiments including, Ntsr1-GN209-Cre and Grp-KH288-Cre. For optogenetic tagging experiments, Ai32 mice (JAX: 012569) expressing channelrhodopsin-2 in Cre-dependent manner, were crossed with Ntsr1-GN209-Cre and GRP-KH288-Cre. Mice were kept on a 12:12 h light:dark cycle and sterilized food pellets and water were provided *ad libitum*. Experiments were performed during the light phase.

## METHOD DETAILS

### Surgical procedures and viral injections

#### General surgical procedures

Anesthesia was achieved using an i.p. dose of ketamine (100 mg/kg) paired with medetomidine (1 mg/kg). Deeply anesthetized mice, verified by a negative pedal reflex, were secured in a stereotaxic frame (Narishige, SR-5N) atop a homeothermic heating pad. Prior to incision, eyes were coated with Dura tear ointment (Novartis, 288/28062–7), and the surgical site was prepped by shaving and applying iso-betadine. To provide local anesthesia, 0.5% lidocaine (0.007 mg/g) was injected subcutaneously into the scalp, after which the skull was exposed via a standard midline incision and tissue retraction.

#### Viral injections

After exposing the skull, three holes were made bilaterally above the superior colliculus within a 1 mm^2^ field anterior of lambda and starting at the midline. Holes were made by gently rotating a needle. For stereotaxic injections, we used glass capillary micropipettes (Wiretrol II, Drummond Scientific, 5-000-2005) pulled to a tip size of roughly 30 µm using a laser puller (P-2000, Sutter Instrument). The pipettes were controlled by an oil hydraulic micromanipulator (MO-10, Narishige). We injected a total volume of 200 nL per hole, divided into two steps. The virus pAAV8-hSyn-DIO-hM4D(Gi)-mCherry (Addgene plasmid #44362) was used for chemogenetic inhibition, while AAV2-hSyn-EYFP (UNC vector core, AV4376E) and pAAV8-hSyn-DIO-mCherry (Addgene plasmid #50459) were used for control experiments.

After the injections, the skin was glued closed using Vetbond tissue adhesive (3M, 1469). Post-operative care included immediate i.p. administration of a painkiller (buprenorphine, 0.2 mg/kg) and an antibiotic (cefazolin, 15 mg/kg), followed by recovery on a heating pad. Over the following 3 days, animals were monitored closely and provided with soft food and antibiotic-treated water (emdotrim, Ecuphar, BE-V23552). Extra doses of buprenorphine and cefazolin were given if needed based on the animal’s recovery.

#### Electrophysiology recordings

Following skull exposure, a metal headpost was fixated to the skull using dental cement (Superbond C&B, Prestige-dental). Craniotomies were performed exclusively over the left hemisphere, targeting either the superior colliculus (‘colliculus’) or the lateral posterior thalamic nucleus (LP). To access the colliculus, a ~1 mm^2^ rectangular craniotomy was made relative to lambda with center: AP 0.5, ML 0.5. To target the LP, a linear craniotomy was made relative to bregma, spanning from AP −2.1, ML 1.6 to AP −3.1, ML 2.1.

For acute experiments, craniotomies were performed 1–2 days prior to the recording session. For chronic implantations, Neuropixels 2.0 probes (imec) were implanted immediately following the craniotomy.

Prior to insertion, the probe shank was coated with a fluorescent dye (DiD, ThermoFisher) to allow for post-hoc tract reconstruction, and the base was covered with a mixture of bone wax and mineral oil. The probe was then slowly inserted to a target depth of ~3500 µm at a rate of ~4 µm/s (taking approximately 30 minutes). Once the target depth was reached, the probe was attached to the skull using Super-bond C&B dental cement.

#### Optic fiber implantations

Optic fibers (200 μm diameter, 3.5 mm long; Doric Lenses) were implanted contralaterally to prevent spatial overlap with the recording probes. Optic fibers were inserted from the right hemisphere (AP +0.6, ML +1.5) at an elevation angle of 56° and 54° to target narrow- and wide-field cells in the left hemisphere, respectively. The fiber was lowered 1.8 mm to place the tip at ML 0, DV −1.1 to −1.2. The entire implant was then fixed in place using dental cement (Superbond C&B, Prestige-dental).

#### Cranial windows for fUSI

Following anesthesia and exposure of the skull, the lateral and posterior muscles were carefully retracted, and veterinary glue (vetbond) was applied to the exposed tissues. Then a large bilateral craniotomy spanning from Bregma +1.5mm to Bregma −5.5 was made by drilling ~1mm2 plaques and carefully removing them with thin tweezers. Once the skull had been removed we cemented a 3D printed headplate to the remaining skull on prefrontal areas, sides and back of the head using dental cement. Finally, the cranial window was covered with silicone elastomer. Mice were allowed to recover for at least 7 days before starting handling and habituation protocols and were given painkillers (Buprenorphine 0.2 mg/kg I.P.), antibiotics (Cefazolin 15 mg/kg I.P.) and anti-inflammatory (Dexamethasone 0.1 mg/kg) drugs for 5 days.

### Histology

Brains were extracted and post-fixed overnight using 4% paraformaldehyde (HistoFix, Roche). Vibratome sections (100 µm) were collected in 1x PBS and were incubated in blocking buffer (1x PBS, 0.3% Triton X-100, 10% Donkey serum) at room temperature for 1 hour. To enhance native fluorophore signal and counterstain nuclei, sections were incubated for 48 h at 4°C with dye-conjugated single-domain antibody fragments (nanobodies) and DAPI in staining buffer (1× PBS, 0.3% Triton X-100, 3% normal donkey serum): ChromoTek GFP-Booster Alexa Fluor 488 (gb2AF488, 1:1,000) for ChR2-GFP, ChromoTek RFP-Booster Alexa Fluor 568 (rb2AF568, 1:1,000) for hM4Di-mCherry, and DAPI (Roche, 10236276001, 1:1000). Sections were then washed three times for 10 min in 1x PBS with 0.3% TritonX-100 and once in 1x PBS, covered with mounting medium (Dako, C0563) and a glass coverslip. Confocal microscopy was performed on a Zeiss LSM 710 microscope. Images of areas with Chr2- and hMD4-expressing cells, the fiber location and the Neuropixels track labelled with DiD were obtained using a 10x (plan-APOCHROMAT 0.45 NA, Zeiss) objective.

### Freely-moving behavior experiments

#### Behavior setup

Visually guided freely-moving behavior assays were performed in a custom-made setup (L: 34 cm, W: 80 cm, H: 50 cm). The floor was made of transparent, red-tinted acrylic to enable video recordings from the bottom. The red-tinted acrylic shelter (height: 10 cm, length: 24 cm, width: 11 cm) was placed in one corner. The ceiling of the setup consisted of visual display (LG 34UM69G-B, 34” diagonal Class 21:9, 250 cd/m^2^ mean luminance, 60 Hz refresh rate) covered with three logarithmic filters to provide moon-light conditions (0.3 lux) and a matt film to prevent reflections. The setup was illuminated with an IR LED strip mounted inside a diffusive plastic enclosure and placed beneath the arena’s floor on the edges.

Behavior was recorded with a near-IR camera placed at the bottom (BASLER acA 1300-60g mNIR) at 60 frames per second with a fixed 6 mm focal lens (Ricoh Lens FL-CC0614A-2M). The camera was triggered with a custom-made Bonsai workflow (Lopes et al., 2021, 2015).

#### Behavioral procedure

All the mice used in the experiments were tested prior to injection of the virus to examine their native responsiveness to stimuli. Mice were habituated to moonlight conditions for 20 min and familiarized with the setup for 10 min before the test. The test included exposure to a set of threatening visual stimuli that were manually triggered using custom-made Bonsai (Lopes et al., 2015) workflow. Mice were exposed to a maximum of 5 repetitions of each stimulus. Mice that did not respond robustly to all stimuli were excluded from the study.

Stimuli used in the pretest were the following: 4° sweeping dot, 20° sweeping dot and black looming (full-contrast dot expanding linearly from 2° to 50°).

#### Stimuli

Visual stimuli were created using the BonVision (Lopes et al., 2021) package within Bonsai. The appearance of the stimulus was registered with a photodiode (Gravity: Analog Grayscale Sensor for Arduino; DFRobot, SKU:DFR0022) placed near the edge of the screen. The animal’s position was tracked in real-time and stimuli were automatically presented when the animal entered an ROI in the middle of the setup after a random inter-trial interval between 30 to 60 s. All stimuli were presented on a gray background and were gamma corrected to compensate for the non-linear luminance effect of the display. We presented looming stimuli of different contrasts: black, gray, and white looming (−98%, −30%, and 111% Weber contrast). Looming stimuli expanded linearly from 2° to 50° in 250 ms. After the expansion phase, the stimulus stayed on the screen in its final size on the screen for another 250 ms. Dimming stimuli consisted of a static 50° disc that linearly dimmed in luminance from the background gray to black (–98% Weber contrast) over 250 ms, followed by a 250 ms hold at maximum contrast. Each trial consisted of three consecutive stimulus presentations with 500 ms inter-stimulus interval displaying background. Stimuli were presented in random order and repeated 5 trials.

### Head-fixed electrophysiology

#### Head-fixed recording setup

Mice were head-fixed in the center of a custom-built 50-cm diameter panoramic display. We displayed visual stimuli using a Texas Instruments projector (DLPDLCR3010EVM-G2; 1280 × 720 resolution, 60 Hz refresh rate, 5 cd/m^2^ average luminance) coupled with a Canon condenser (EF 50 mm F1.4 USM) and a Peleng fisheye lens (8 mm f3.5 M42). A photodiode mounted directly on the lens tracked the exact stimulus timing. Neuropixels data were sampled at 30 kHz via an imec headstage (HS_1000) and PXIe acquisition module (PXIE_1000) housed in an NI chassis (PXIe 1071). We recorded the photodiode using a PXIe I/O module (PXIe-6341, 30488 Hz sampling rate). To temporally align the data streams, an Arduino generated a master synchronization pulse that was sent simultaneously to the Neuropixels and I/O modules.

#### Head-fixed recording protocol

Mice were habituated to the recording rig for 1–2 days, beginning at least 3 days after headposting. On recording days, mice were head-fixed atop a spherical treadmill centered within the panoramic display.

For acute recordings, Neuropixels 1.0 or 2.0 probes (imec) were coated with a fluorescent tracker (DiD, DiI, or DiO; Thermo Fisher) and lowered at ~4 µm/s until the tip rested 0.5–1.5 mm beyond the target structure. The exposed tissue was covered with either saline or artificial cerebrospinal fluid (aCSF: 150 mM NaCl, 5 mM K, 10 mM D-glucose, 2 mM NaH_2_PO_4_, 2.5 mM CaCl_2_, 1 mM MgCl_2_, 10 mM HEPES; pH adjusted to 7.4 with NaOH). We then waited 30 minutes for the brain tissue to settle before initiating the recording.

Electrophysiological data were sampled from 384 channels distributed over 3840 μm (across 1–2 shanks), chosen to increase channel density within the region of interest.

#### Stimuli

All visual stimuli were presented using custom OpenGL-based software. To quickly estimate receptive field locations online, we presented a 4° white square sweeping across the right hemifield at azimuths from 0° to 90° in 15° increments. For recordings from the colliculus, we centered all subsequent stimuli at the azimuth and elevation that elicited the strongest visual response in the superficial layers. For lateral posterior nucleus (LP) recordings, the stimulus location was fixed at either 30° azimuth/30° elevation or 45° azimuth/45° elevation.

Looming stimuli of different contrasts were presented on a uniform gray background (background illuminance: 19.3 lx; black: −96%, −77%, and −41% Weber contrast; and white: 145% and 280% Weber contrast). Standard trials consisted of three consecutive expanding disks. During each presentation, the disk expanded from 2° to 50° in 250 ms, remained at maximum size for 250 ms, and was followed by a 330 ms inter-stimulus background. Trials were separated by a randomized 7–15 s inter-trial interval. The dimming stimulus linearly changed from background contrast to −96% negative Weber contrast over 250 ms.

#### Optogenetic tagging

Following visual stimulation, we performed optogenetic tagging via the implanted optic fiber. We delivered 500 light pulses (30 ms duration) separated by uniformly randomly sampled inter-pulse intervals ranging from 100 ms to 1 s. Pulse edges were shaped with 1 ms on and off ramps generated by an Arduino. We empirically titrated the optical power for each recording based on online neural responses observed in SpikeGLX. Testing initially began at 0.150 mW; if no driven activity was detected, we increased the power to 0.185 mW, whereas visible evoked responses prompted a reduction to 0.100 mW or 0.050 mW.

### Freely-moving electrophysiology

#### Freely-moving recording setup

Recordings were conducted in a modified behavior setup (L:19, W:80 cm). The reduced setup width prevented cable torque on the Neuropixels cable. To prevent tangling of the Neuropixels cable, we used motor-assisted electrical rotary joint (doric; AERJ_24_HDMI+4). Neuropixels data were sampled at 30 kHz via an imec headstage (HS_1000) and PXIe acquisition module (PXIE_1000) housed in an NI chassis (PXIe 1071). We recorded the photodiode signal using a PXIe I/O module (PXIe-6341, 30,488 Hz sampling rate). To temporally align the Neuropixels and I/O data streams, the PXIe acquisition module generated a master synchronization pulse that was sent to the I/O modules. Alignment of the camera frames and the photodiode was achieved by logging each entry of the photodiode and camera frame in Bonsai.

Visual stimuli were presented on an overhead monitor and a lateral monitor. Unlike the behavioral setup described above, we omitted the neutral density filters (0.3 lux) and matte diffusing film to maximize stimulus salience.

#### Freely-moving recording protocol

Mice were habituated to the recording rig for 2 days, beginning at least 5 days after Neuropixels implantation. In each habituation session, mice were free to explore the arena for 15 minutes. They were then moved to the head-fixed recording setup followed by 15 minutes of head-fixation. On recording days, mice performed the freely moving task first followed by head-fixed recordings with the head-fixed experimental protocols described above.

#### Stimuli

Visual stimuli were presented on either the overhead or lateral monitor based on the receptive field location of the recorded superior colliculus cells. Stimulus parameters (black looming, white looming, and dimming) were identical to those described above (see “Freely-moving behavior experiments”). Visual stimuli were manually triggered when animal ran across the middle of the setup.

### Functional ultrasound imaging

#### fUSI setup

For fUSI experiments, the same dome was used as for the head-fixed electrophysiology. In these experiments, the dome was combined with a volumetric fUSI acquisition system (Brunner et al., 2020). Briefly, here we used a workstation controlling a Vantage 256-channel scanner system ((Vantage 256, Verasonics, USA), coupled to a 2D-matrix transducer of 32 × 32 (1024) piezoelectric elements (MAT 15.0/32×32-SiX, Vermont), where each element has a size of 0.3 × 0.3 mm2, the total size of the potential imaging area is 9.8 × 9.8 mm2. Each trial was triggered and saved separately using the fUSI acquisition start and end as the master trigger for starting visual presentations and both starting and ending camera recordings.

#### Habituation and fUSI imaging preparation

After at least 7 days after surgery, mice were habituated to handling for 2 days followed by increasing periods of head fixation on the air-cushioned ball system for at least 4 days. Starting with 15min and reaching 1h in ~4 days. Once habituated, before an imaging session, the silicone cap was removed, the brain was covered with aCSF and allowed to rest for 5 min, then it was removed and quickly covered with 3% agarose solution. Once fully solidified, the agarose was covered with ultrasonic gel and the ultrasound probe was lowered until being ~1mm from the agarose surface and ~3mm from the brain surface. Following, we performed the functional imaging acquiring 70 whole-brain volumes of Doppler images per trial at 0.5 frames per second.

#### fUSI aquisition sequence

Functional ultrasound acquisitions followed the methodology described previously (Brunner et al., 2020). Using this approach, we imaged a brain volume of 9.6 × 8.1 × 8 mm^3^(lateral, anteroposterior, and dorsoventral dimensions, respectively), with a spatial resolution of 150 × 150 × 100 µm^3^.

#### High-quality vascular imaging

At the end of each session, we acquired a high-quality anatomical vascular image. This image was generated using the SUSHI method (Bar-Zion et al., 2018), with a 3D point-spread function instead of the 2D point-spread function used in the original method.

The acquisition sequence consisted of 12 angles, defined by lateral and anteroposterior steering angles, respectively: (−9°, 0°), (−6°, 0°), (−3°, 0°), (3°, 0°), (6°, 0°), (9°, 0°), (0°, −9°), (0°, −6°), (0°, −3°), (0°, 3°), (0°, 6°), and (0°, 9°). Beamforming was performed using isotropic voxels with a size of 100 µm.

We averaged 100 µDoppler spectra and divided the resulting spectrum into 25 frequency bands. Each band was super-resolved using isotropic 50-µm voxels and the FISTA algorithm (ref. FISTA). The final high-quality vascular image was obtained by averaging the 25 super-resolved images.

#### Stimuli

For fUS imaging experiments, mice were placed in a panoramic set up as for the head-fixed electrophysiology recordings. Here mice were presented with a black looming disc, a dimming disc, and a large sweeping disc. Here the disk expanded from 2° to 50° in 330ms, and remained for 500ms. Each trial consisted of 5 consecutive expanding disks with a 170ms inter-stimulus interval. Similarly the dimming stimulus had a 50° diameter and consisted of 5 repetitions per trial, using the same temporal structure. Finally, the sweeping disc was also 50° in diameter, appeared gradually from behind a mask of background color, and was only visible within the 50° of visual space that it crossed at a speed of 25°/s. A trial consisted of 2 sweeping stimuli with a 0ms inter-stimulus interval. Each stimulus was presented in 20 trials per session in random order. Trials consisted of a fixed 70 fUSI frames that corresponded to 15s baselines, followed by a stimulus (triggered at the onset of frame 30), and ended 20 seconds after stimulus onset. All these stimuli were presented on the right side of the mouse, centered at 45° elevation and 45° azimuth. Sessions were split in looming + dimming (L&D), and looming + sweeping (L&S). In L&D sessions each stimulus was presented at elevations −10° and 45°, 20 trials in each location. For this work, we focused only on the 45° elevation stimuli. In L&S sessions only 45° stimuli were used.

## QUANTIFICATION AND STATISTICAL ANALYSIS

### Analysis of behavioral data

Animal posture and kinematics were extracted from overhead video recordings utilizing DeepLabCut (Mathis et al., 2018). Tracking coordinates with a prediction confidence threshold below 0.9 were discarded and linearly interpolated. Running speed was calculated using the positional coordinates of a marker tracking the approximate center the abdominal region of the mouse. To reduce tracking noise, the resulting velocity traces were smoothed using a 1D Gaussian filter with a standard deviation of ~6 frames (equivalent to ~100 ms). Trials in which the animal’s snout remained inside the shelter for the entirety of the recording were excluded from analysis. Visually-evoked innate behavioral responses were quantified strictly within a 3 s window following stimulus onset. Behaviors were classified computationally based on the following thresholds.

Escape: Body-center speed > 12 cm/s for longer than 166 ms (10 frames). The escape bout needed to be initiated during the stimulus, outside the shelter and terminated within the shelter.

Immobility: Body-center speed < 8 cm/s sustained for longer than 166 ms (10 frames).

To evaluate the effect of chemogenetic manipulation (CNO versus control) on visually-evoked behaviors while accounting for intermouse variability, we fitted a binomial generalized linear mixed-effects model (GLME) using pymer4. Escape behavior was modeled as a binary outcome. The model incorporated the experimental condition, trial number, and pre-stimulus onset speed (defined as the mean speed from −0.5 to 0 s relative to stimulus onset) as fixed effects. To account for repeated measures, mouse identity was included as a random intercept (Formula: Escape ~ Condition + Trial + Onset_speed + (1 | Mouse_ID)) (Fig. S1).

### Analysis of Neuropixels recordings

#### Spike sorting

The recorded data (SpikeGLX v20230101-phase30) was preprocessed using SpikeInterface v0.102.3 (Buccino et al., 2020). For optotagging experiments, optical stimulation artifacts were removed prior to preprocessing via linear interpolation of the raw voltage traces across 1-ms windows (−0.5 to +0.5 ms relative to stimulation onset and offset). All recordings were then preprocessed following the International Brain Laboratory destriping pipeline (Laboratory, 2024) to remove spatially correlated noise. Spike sorting was performed using Kilosort 3 for head-fixed experiments and Kilosort 4 for freely moving experiments (Pachitariu et al., 2024, 2016).

Sessions where animals transitioned immediately from the freely moving open-field setup to the head-fixed dome setup were temporally concatenated and spike-sorted as single continuous recordings. Putative single units were curated using metrics from the Allen Institute ecephys pipeline (https://github.com/alleninstitute/ecephys_spike_sorting). Clusters were included in single unit analyses if they met the following criteria: amplitude cutoff < 0.1, overall firing rate > 0.5 Hz, and presence ratio > 0.6. Our criteria for the number of inter-spike interval (ISI) violations depended on the number of spikes of the unit: units with >2,000 spikes required an ISI violation score < 0.5, while units with 500–2,000 spikes required a refractory period violation ratio < 0.005.

#### Detection of visually responsive units

Single unit responsiveness to visual stimuli was determined using the parameter-free Zeta test (Montijn et al., 2021). Spike trains were aligned to stimulus onset, with trial epochs extracted from −2 to +8s. The statistical significance of the stimulus-evoked response was evaluated over a response window from 0 to 4s post-stimulus. We resampled 500 times to generate the null distribution. A unit was considered responsive to a visual stimulus if the ZETA test yielded a significance level of p<0.05. Units were classified as visually responsive is they met this criterion for at least one of the presented stimuli.

#### Firing rate estimation

Continuous firing rates were estimated by convolving spike trains with a non-causal Gaussian kernel (σ = 25ms) using the Elephant package (Denker et al., 2026).

#### Visual tuning and lifetime sparsity

For each visually responsive unit, estimated firing rates were averaged across trials. Baseline activity, defined as the mean firing rate from −1.25 to −0.25 s relative to stimulus onset, was subtracted from these trial-averaged traces. The 250ms directly preceding the stimulus onset was excluded to prevent backward data leakage from the non-causal smoothing kernel. To construct visual tuning curves (Fig 2H,K), we extracted the peak evoked firing rate for each stimulus. The peak firing rate was defined as the 98^th^ percentile (to ensure robustness against outliers) during the first 0.5s after stimulus onset. This window includes both the dynamic and static phases of the first stimulus presentation. For each unit, tuning curves were then normalized by dividing the peak stimulus responses by the maximum peak response across all stimuli.

Lifetime sparsity was quantified using the adjusted Treves-Rolls sparsity index, defined as:

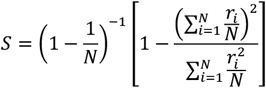

where *N* is the total number of stimuli and *r*_*i*_ is the peak evoked firing rate for stimulus *i* (Vinje and Gallant, 2000).

#### Detection analysis

To evaluate the ability of neural population to detect the presented stimuli, we trained a logistic regression classifier on the population activity to discriminate baseline from stimulus-evoked activity. Analysis was performed on putatively single and visually-responsive units. For each unit, baseline firing rate was calculated as the mean activity during the pre-stimulus window (−1.25 to −0.25s relative to stimulus onset). Stimulus-evoked firing rate was calculated as the mean activity during the first 0.5s of the stimulus (includes both the expansion and static phases of the first stimulus presentation). The logistic regression classifier was implemented using scikit-learn. Model performance was validated using repeated stratified k-fold cross-validation (k=5; RepeatedStratifiedKFold) to preserve balanced class distributions across splits.

Model discriminability was evaluated using receiver operating characteristics (ROC) analysis on the probability estimates of the classifier. Predicted posterior probabilities for the stimulus class were computed on test trials using the predict_proba function in scikit-learn. The area under the ROC curve (ROC-AUC) was computed across folds to quantify separability independently of decision threshold.

#### Dimensionality estimation using Participation Ratio

To quantify the dimensionality of the neural representations in the dSC and LP, we calculated the participation ratio 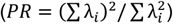 of the trial-averaged firing rates, where λ_*i*_ are the eigenvalues of the neural covariance matrix. Because PR is highly sensitive to the number of neurons (N), we controlled for unequal yields between the dSC (N = 479) and LP (N = 441) pseudo-populations via a strict subsampling protocol. For each region, we randomly subsampled 100 neurons without replacement and computed the PR across all visual stimuli. This process was repeated 1,000 times to generate a mean PR and 95% confidence intervals. To assess statistical significance, we utilized a non-parametric two-tailed permutation test. Briefly, all neurons from both the dSC and LP were pooled into a single pseudo-population. In each of 10,000 permutations, region labels were randomly shuffled, and 100 neurons were drawn (without replacement) for each surrogate group to calculate a null difference in PR. The pvalue was defined as the proportion of permutations where the absolute null difference exceeded the absolute observed difference.

#### Decoding analysis – pseudopopulation

To assess whether high-threat stimuli could be discriminated from low-threat stimuli using population activity, we trained support vector classifiers (LinearSVC, Scikit-learn) with a linear kernel and balanced class weighting. We used l2-regularization (C=0.1). Classifiers were trained to discriminate binary groupings of visual stimuli: the threat category consisted of black looming trials; the non-threat category consisted of trials of gray looming, white looming, and dimming stimuli.

To preprocess the data, the activity of each cell was normalized by subtracting the baseline for each trial (−1.25 to −0.25 s relative to stimulus onset and dividing by the standard deviation of the concatenated stimuli. To reduce the contribution of rapid changes associated with habituation to the visual stimuli, we excluded the first two trials of each stimulus from our analysis.

Cross-validation was performed using a Leave-One-Out (LOO) scheme across physical stimulus presentations (N = 8 trials). To decouple trial-to-trial noise correlations while preserving individual neuronal response statistics, training datasets were constructed via pseudo-trial resampling. On each cross-validation fold, the n-1 physical training trials were sampled with replacement independently for each neuron to generate N = 100 synthetic pseudo-trials per stimulus condition. The held-out physical trial was reserved exclusively used for out-of-fold testing. Model performance was quantified using balanced decoding accuracy to eliminate bias arising from unequal stimulus class sample sizes.

To evaluate decoding performance and population variance over time, we fitted PCA on the concatenated visual responses across stimuli and trained a separate linear classifier for each 5 ms bin (after smoothing with a 25 ms std gaussian kernel). To control for population size differences across brain regions, a random subpopulation of N = 100 visually responsive neurons was drawn without replacement on each bootstrap iteration.

Dimensionality reduction was performed within each cross-validation fold to prevent overfitting and extract low-dimensional signal dynamics: Peri-stimulus time histograms (PSTHs) were computed from the training pseudo-trials across all stimulus conditions and concatenated across time. Principal Component Analysis (PCA; 10 principal components) was fitted on the concatenated training PSTHs. Both the training pseudo-trials and held-out physical test trials were centered using the training mean and projected into the fitted PCA subspace. A separate linear classifier was fitted independently at each 5-ms time bin using the low-dimensional training representations. Balanced decoding accuracy was evaluated on the held-out test trial and averaged across LOO folds. To establish empirical chance performance, a shuffle null distribution was generated at each time bin by permuting stimulus class labels across 10 shuffle iterations per fold.

To determine whether the population readout dimension aligned with dominant shared variance in the neural latent space, the proportion of signal variance captured by the decoder was quantified across time. Specifically, the variance explained by the decoder was calculated using its unit-normalized weight vector 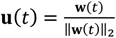 principal component variance ratios *λ*_*i*_ derived from the training covariance matrix:

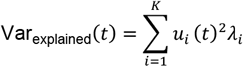

Signal variance was summarized within a target post-onset window (25–75 ms) and visualized as bootstrap means accompanied by nonparametric 95% bootstrap confidence intervals (Fig. 4H). Within-region significance was calculated by generating a null distribution across 10 stimulus-label shuffles per fold and computing the proportion of null iterations exceeding observed variance. Between-region differences were evaluated by computing the empirical two-tailed pvalue directly from the bootstrap difference distribution (Var_*A*_ − Var_*B*_). To quantify decoding efficiency as a function of the number of cells (Fig. S3D), we trained the classifiers directly on the preprocessed neural firing rates. Single-trial firing rates were averaged across the 25–75 ms post-onset window prior to classification. Subpopulations of varying sizes (*N ∈* {1,10,25,50} neurons) were randomly drawn without replacement across 100 bootstrap iterations to plot balanced accuracy (mean ± s.d.) as a function of population size.

#### Population activity in freely-moving animals

To assess population-level responses during behavior, we pooled single-unit and multi-unit activity.

We defined escape behaviors based on the animal’s running speed. A trial was classified as an escape if the running speed exceeded 25 cm/s and stopped running in a corner of the setup. We visually inspected the automated detections for all sessions. Sessions with automated detection failures were manually annotated. Escape onset was then determined retrospectively from the detected bout by traversing the running speed trace backward in time and marked onset at the first frame where speed fell below 5 cm/s or acceleration was ≤ 0 cm/s^2^.

We used a Receiver Operating Characteristic (ROC) analysis to distinguish escape from non-escape trials based on the neural activity in the dSC and LP (based on Evans *et al*., 2018). The input feature was the cumulative sum of the mean z-scored population activity, extracted from −2 to +1 s relative to escape onset. For non-escape trials, we aligned the neural data to pseudo-onsets randomly sampled from the latency distribution of escape trials. Statistical significance was evaluated on the 100 ms window immediately preceding escape onset. Null distributions for significance testing were generated by randomly permuting the behavioral outcomes of the trials.

#### Shared latent representations across experiments - multiCCA

Single-unit and multi-unit activity were pooled to assess population-level responses. Sessions containing fewer than 15 units were excluded. Firing rates were baseline-subtracted using the mean activity from a 1 s pre-stimulus window. Trial-averaged responses were extracted for six visual stimuli: 100% black looming, 80% black looming, 40% black looming (gray), 50% white looming, 100% white looming, and dimming. Firing rates were normalized by dividing by the cross-trial standard deviation.

To identify shared latent dynamics across animals, we applied MCCA using the cca_zoo Python package (Chapman and Wang, 2021). Single-session dimensionality was first reduced using PCA, followed by latent space alignment via MCCA on trial-averaged head-fixed data. Cross-session alignment was quantified by computing the average pairwise Pearson correlation across all session pairs for each Canonical Variate. To determine statistical significance, we generated a null distribution using 100 permutations, in which stimuli were shuffled independently for each session prior to refitting the model.

#### Decoding and freely-moving analysis – multiCCA

We trained a linear Support Vector Machine (SVM) on head-fixed data to decode threat versus non-threat stimuli. The 100% and 80% black looming stimuli were classified as threat, while all other stimuli were classified as non-threat. The input features consisted of the mean multiCCA-transformed activity within 25–75 ms post-stimulus onset. The SVM was trained using balanced class weights, and performance was evaluated using leave-one-session-out cross-validation. Decoding accuracy was quantified using balanced accuracy. Significance against chance level (0.5) was assessed using a onesided Wilcoxon signed-rank test.

To evaluate the decoder while the animal was behaving, freely-moving neural data were projected into the head-fixed multiCCA latent space. Single-trial freely-moving data were transformed using the pre-fit MCCA weights, z-scored, and smoothed with a 250ms bin moving average. The pre-trained head-fixed SVM was then applied to the freely-moving latents to extract decision values across time.

Escape bouts were classified based on running speed and corner distance (See methods section ‘Population activity in freely-moving animals’). Escape bouts were predicted using the peak SVM decision value within a 3-second post-stimulus window. These peak values were used to generate a Receiver Operating Characteristic (ROC) curve.

#### Optogenetic tagging

Visually responsive units (we included both single and multi-unit activity) were manually curated to identify which ones were activated by optogenetic stimulation. We subdivided the optotagged units based on their median response onset latencies. Units with a response probability ≥ 10% (of trials) and a response onset < 9 ms were classified as putative secondary cells. Units with latencies greater than 9 ms were classified as non-secondary (other) optotagged cells.

We evaluated the stimulus representations of optotagged populations using Principal Component Analysis (PCA) on trial-averaged firing rates. To compare populations with unequal unit counts, we bootstrapped the PCA. We randomly subsampled each group to 30 cells across 10,000 iterations. This allowed us to estimate the cumulative variance explained and the specific variance captured by the first three principal components. Statistical significance was derived from the 95% confidence intervals of the difference between bootstrapped distributions.

We trained a linear decoder on trial-level pseudopopulation firing rates. The decoder discriminated between a high-threat stimulus (black loom) and lower-threat stimuli (gray loom, white loom, and dimming). We used two randomization tests (1,000 iterations each) to evaluate the contribution of secondary cells:

Sufficiency: We calculated decoding accuracy and variance explained using only secondary cells. We compared this to a null distribution derived from size-matched random subsets of non-secondary dSC cells.

Necessity (Ablation): We calculated decoding performance on the full dSC population excluding secondary cells. We compared this to a null distribution formed by excluding a size-matched random subset of non-secondary cells.

P-values were calculated based on the proportion of the null distribution that equaled or exceeded the metrics observed in the targeted subsets.

#### Modeling of downstream integration of threat

To evaluate how dSC and LP threat representations coordinate escapes, we modeled a two-dimensional state space defined by the threat decoder decision values in the dSC (x-axis) and the LP (yaxis). Stimulus responses were modeled as 2D Gaussian probability distributions. We parametrized the threat and non-threat distributions as *µ*_*threat*_ = (2,2) and *µ*_*non*−*threat*_ = (−2, −2) and *σ*_*threat*_ = *σ*_*non*−*threat*_ = 0.8.

We tested three downstream integration architectures: Competition, Linear Summation, and a non-linear Driver-Gate. We defined the neural drive for escape (*Z*) for each model as follows. The Competition model used subtractive integration, defined as *Z* = *w*_1_*x* − *w*_2_*y* + *b*. The Linear Summation model used additive integration, defined as *Z* = *w*_1_*x* + *w*_2_*y* + *b*. The Driver-Gate model used multiplicative integration, defined as 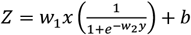 The Driver-Gate bias term was set to *b* = −0.1.

To evaluate how these model parameters influence behavioral predictions, we systematically swept the slope and offset variables across simulations. The varying slope corresponded exclusively to the LP input parameter (*w*_2_) across all models: acting as the LP weight in the Competition and Linear Summation models, and as the LP sigmoidal steepness in the Driver-Gate model. The varying offset corresponded directly to the mathematical bias term (*b*) across all architectures (Fig. S8A). During these sweeps, the dSC weight multiplier (*w*_1_) was held constant.

We simulated neural manipulations to test model validity against *in vivo* behavioral constraints. Manipulations were modeled as linear vector trajectories across the state space. Inhibition trajectories originated from the threat state at (2,2). The dSC inhibition target was (−2,2), and the LP inhibition target was (2, −2). Optogenetic stimulation trajectories originated from the non-threat state at (−2, −2). The dSC stimulation target was (3, −2), and the LP stimulation target was (−2,3). We calculated the simulated escape probability using volume-over-threshold integration. A motor decision boundary separated high-threat from low-threat stimuli where *Z* > 0. We quantified escape probability as the exact volume of the 2D Gaussian distribution exceeding this threshold.

### Analysis of fUSI data

#### Registration

The registration procedure has been described previously (Brunner et al., 2021). Briefly, for each session, the high-quality µDoppler image was registered to the Allen Mouse Brain Common Coordinate Framework (Wang et al., 2020) using an affine transformation. Before applying the affine transformation, all images acquired during the session were interpolated to isotropic 50-µm voxels. The final output consisted of a set of 3D images with isotropic 50-µm voxels, spatially aligned to the atlas.

#### Segmented temporal responses

The registered images were segmented into 318 brain regions per hemisphere defined by the CCF atlas (see the list of regions in the Supplementary Material). For each image, the mean intensity of all voxels within each brain region was calculated. This signal was normalized to its baseline value, defined as the 3 seconds preceding stimulus onset.

Finally, we computed the median for all the trials of the same animal and stimulus condition. This procedure yielded a representative temporal response for each brain region, animal, and stimulus condition normalized by the baseline.

### Statistical Analysis of fUSI Responses

To evaluate the functional ultrasound imaging (fUSI) responses, we analyzed data from over 300 distinct brain regions (see Supplemental Table S1). To quantify the magnitude of the activation in each brain region, we calculated Cohen’s *d* as the primary measure of effect size. We utilized a data-driven bootstrapping approach to determine the reliability of these effects. For each brain region, the observed data was resampled with replacement 5000 times to generate a 95% confidence interval around the effect size. To determine statistical significance, we employed a non-parametric permutation test rather. For each region, we tested the hypothesis that the mean response was greater than baseline. The data points were shuffled 99,999 times to build a null distribution, allowing us to calculate a raw p-value. Because we tested hundreds of brain regions simultaneously, we adjusted the raw *p*-values using the Benjamini-Hochberg False Discovery Rate (FDR) procedure. A region was considered significantly active if its FDR-corrected *p*-value fell below the threshold of 0.05.

### Assessing the Modulatory Effects of CNO

To evaluate how the chemogenetic inhibition of wide-field neurons altered neural responses compared to control (vehicle) conditions, we quantified the difference in activation for each animal. First, we calculated the baseline-subtracted response amplitude for both the CNO and vehicle sessions by taking the mean signal during the stimulus window and subtracting the median baseline activity. Our experimental design was paired within subjects, therefore we directly subtracted the vehicle response from the CNO response to isolate the drug’s effect on each brain region.

These difference scores were then analyzed using the same statistical framework described above, utilizing Cohen’s *d* and 5000 bootstrap iterations to estimate confidence intervals for the effect size. To identify brain regions where CNO caused either a significant enhancement or suppression of the stimulus response, we performed permutation tests (99,999 resamples) on the difference scores. The resulting *p*-values were FDR-corrected for multiple comparisons. Any region exhibiting a statistically significant change (FDR-corrected *p* < 0.05) direction was classified as being modulated by CNO.

## Notes

### Competing Interest Statement

The authors have declared no competing interest.

### Summary of Updates

Missing figure and figure legend. Corrected spelling of authors.

