## Supplemental Figures for "Cooperative processing of visual threat across collicular pathways"

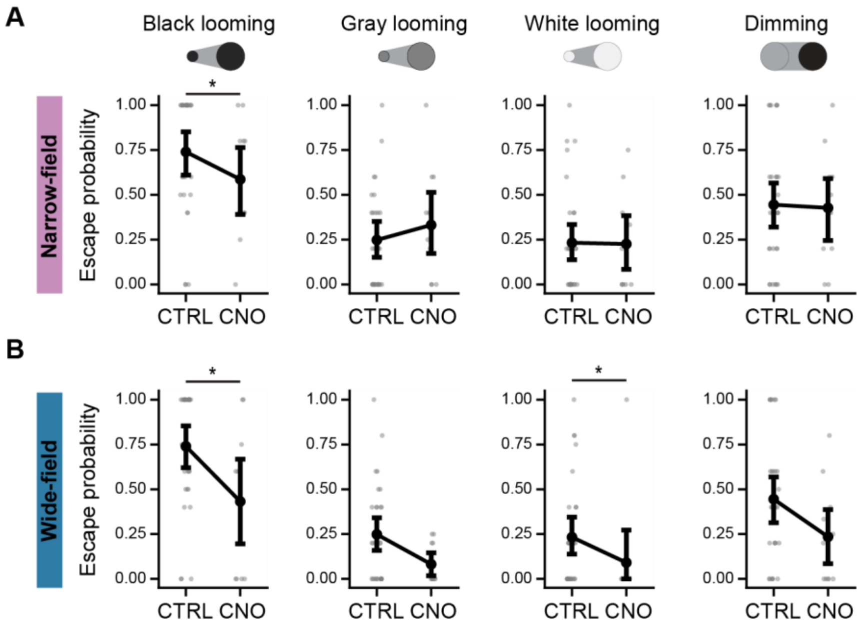


**Figure S1. Effects of inhibition of narrow-field and wide-field neurons on escape behavior. Related to Figure 1. A-B.** Probability of escaping. Individual dots are individual mice (n=11 NF CNO, 11 WF CNO, 17 NF CTRL , 12 WF CTRL mice). Shown is mean and 95% confidence interval. Significance between control (CTRL) and inhibition (CNO) conditions was tested using a binomial generalized linear mixed-effects model (GLME) incorporating the experimental condition, trial number, and pre-stimulus onset speed as fixed effects and mouse identity as random intercept (see methods). Escape behavior was modeled as a binary outcome. *p<0.05 **p<0.01, ***p<0.001.

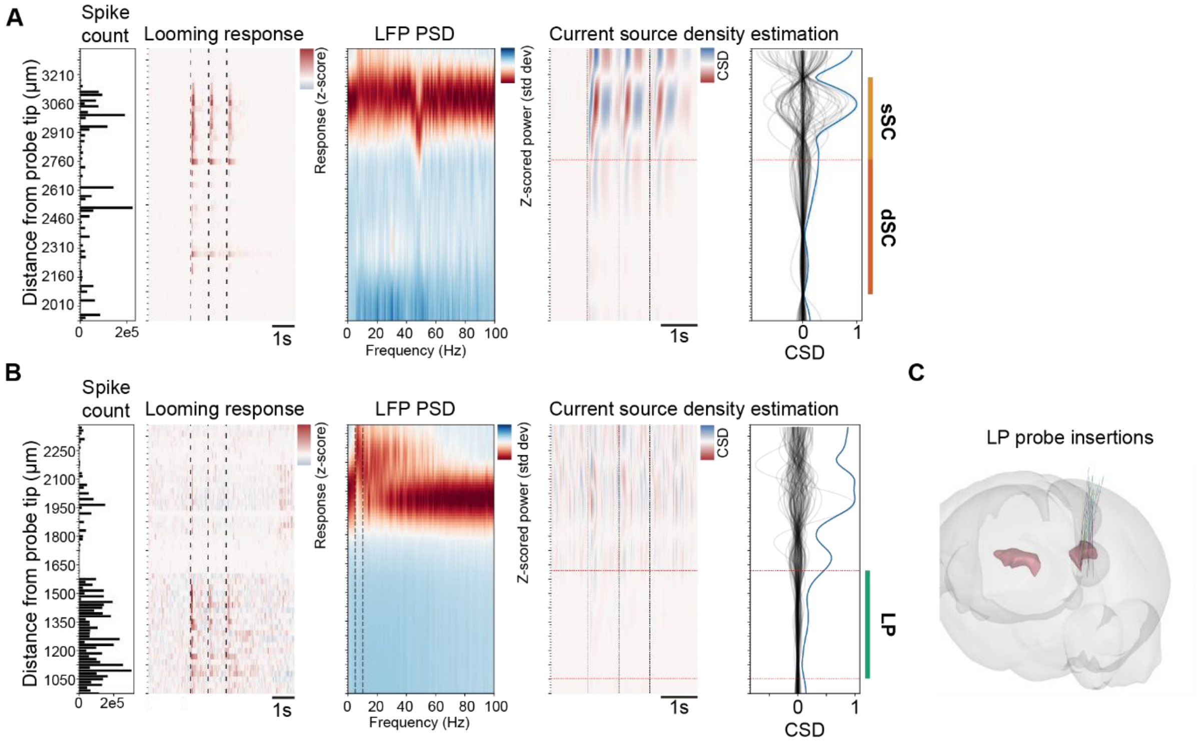


**Figure S2. Region delineation based on physiological markers. A.** The location of the sSC and dSC on the Neuropixels probe were determined based on the zero-crossings in the current source density, visual responsiveness, and histology. The bottom of the sSC was determined by taking the zero crossing and subtracting 100 micrometers. **B.** LP depth was determined using the power spectral density (PSD) of the LFP signals, the spike counts along probe depth, visual responsiveness, and histology. **C.** Visualizations of probe insertions into the LP. These trajectories were determined based on histology.

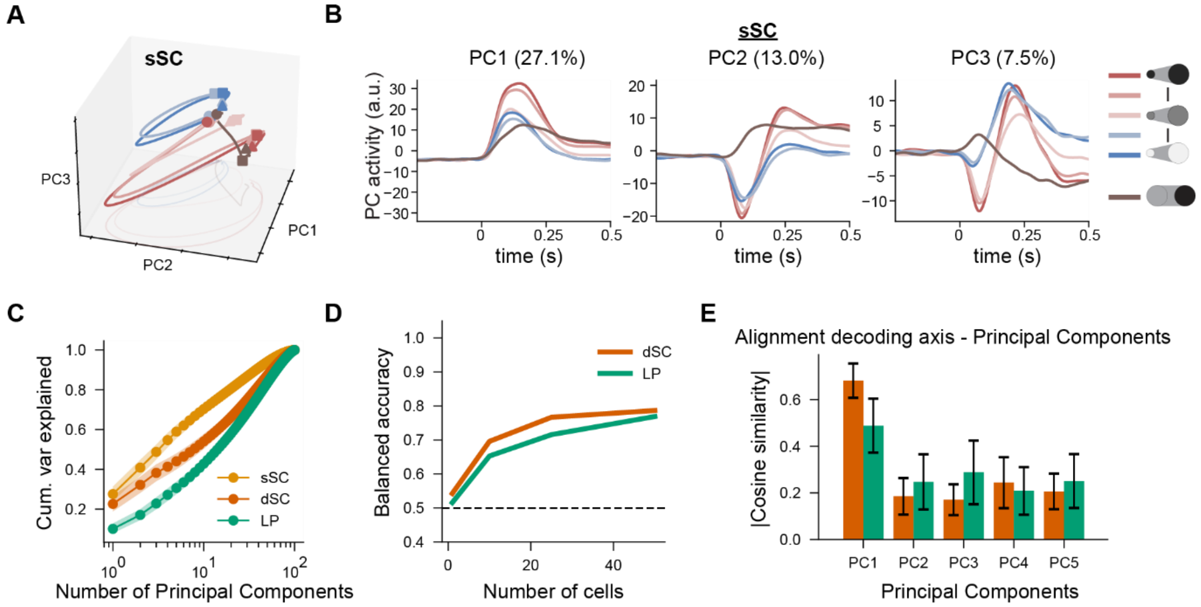


**Figure S3. Low-dimensional representations of threat. Related to Figure 3. A.** First three principal components based on trial-averaged activity of all cells in sSC recordings. **B.** First three principal components across time for the looming stimuli across contrasts and dimming stimulus in the sSC. Looming contrasts: 100% black, 80% black, 40% black (gray looming), 50% white, and 100% white. **C.** Cumulative variance explained in the sSC, dSC, and LP. Median±95% confidence interval (bootstrapping of 100 cells). **D.** Decoding accuracy across number of cells. **E.** Absolute cosine similarity between the decoding axis and the principal components. Mean±std of time window 25-75ms after stimulus onset.

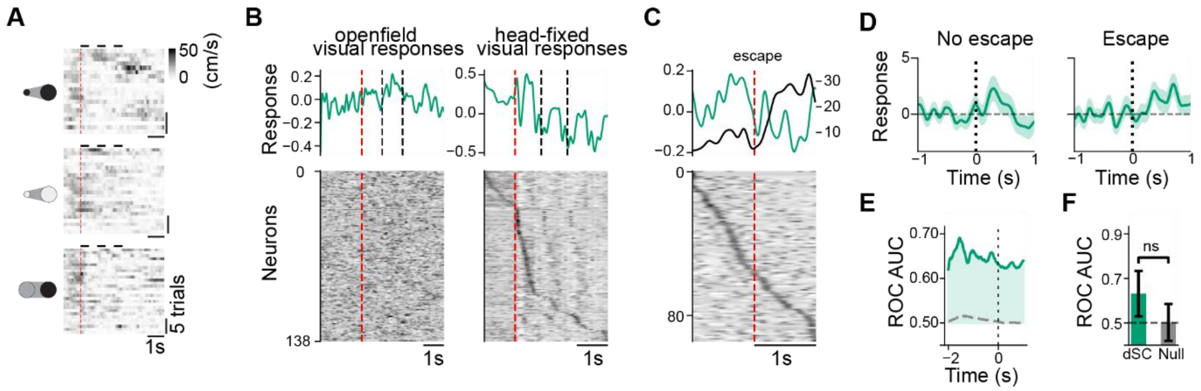


**Figure S4. LP responds to visual stimuli but does not predict escape initiation. Related to Figure 5. A.** Running speed heatmaps across trials aligned to stimulus onset (red dashed line) for black looming, white looming, and dimming stimuli. **B.** Mean z-scored population activity (top) and z-scored response heatmaps (bottom) aligned to visual stimulus onset in open-field (left) and head-fixed (right) setups. Averaged across stimuli. Cells are sorted by peak response latency in the head-fixed setup (N=138 cells, N=2 mice). **C.** Mean z-scored population activity (top) and z-scored response heatmaps (bottom) aligned to escape onset in the open-field setup. Cells are sorted by peak response latency. **D.** Mean LP population response to black looming stimuli on escape trials versus non-escape trials. Zoomed in on first loom repetition indicated by overhead bar. Shaded areas represent mean±SEM across cells. Two-sided Mann-Whitney U test (p=0.867; 20 trials). **E.** Receiver operating characteristic area under the curve (ROC-AUC) over time aligned to escape onset. Gray dashed line denotes chance-level AUC (shuffled null distribution). **F.** Mean ROC-AUC values during the 100ms preceding escape onset. Mean±SD of bootstrapped trials. Permutation test was used to compare the dSC performance against a null distribution (p=0.159; 72 trials).

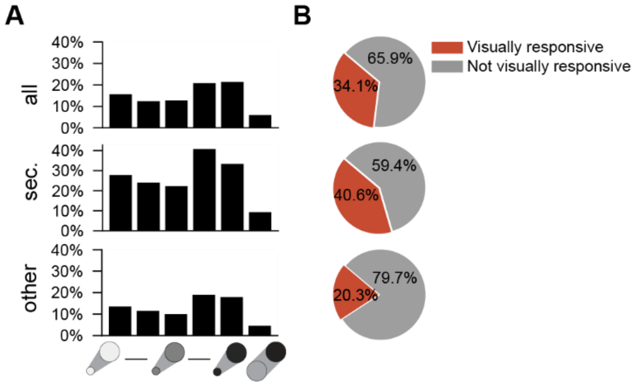


**Figure S5. Proportions of visual responsiveness in optotagged cells. Related to Figure 4. A.** Proportion of cells responsive to distinct visual stimuli across three populations: all recorded cells (n = 1567), putative secondary optotagged cells (sec.; N=64). and other optotagged cells (other; N=123). **B.** Proportion of cells visually responsive to at least one stimulus for each group from A. Orange slices represent the percentage of visually responsive cells; grey slices represent non-responsive cells.

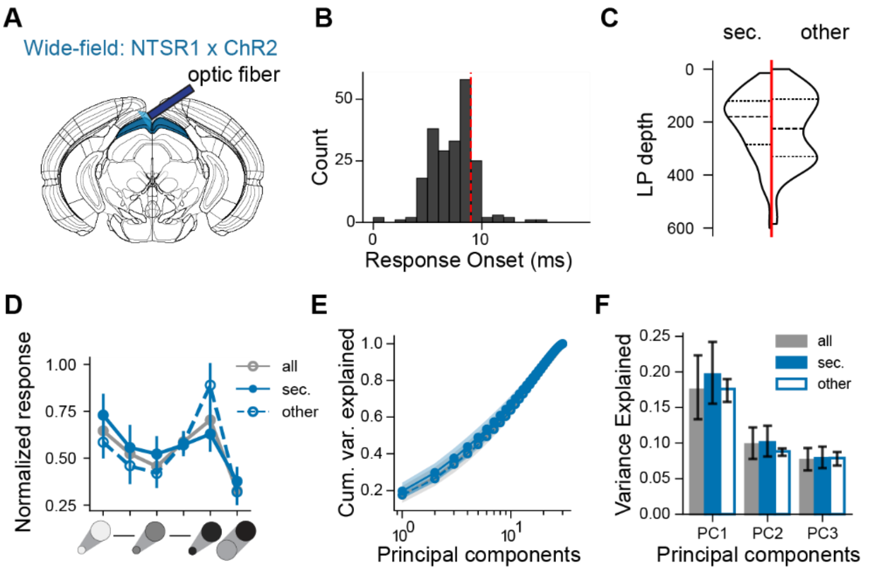


**Figure S6. Wide-field cells target global population in the LP. Related to Figure 4. A.** Schematic of the experimental setup. An optic fiber was implanted in NTSR1xChR2 mice to optogenetically activate wide-field cells while simultaneously recording the activity of LP cells using Neuropixels. **B.** Distribution of response onset latencies to optogenetic stimulation. Cells with a response onset latency <9ms (red dashed line) were classified as putative secondary cells (sec.) (N = 5 mice; $N_{sec}$=172 cells, $N_{other}$=43). **C.** Anatomical distribution of putative secondary and other optotagged cells across LP depth. Depth is measured relative to the surface of the LP. Dashed lines indicate median and IQR. **D.** Normalized stimulus preferences for all visually-responsive LP cells. Responses are normalized to the maximum response across stimuli. Point-and-whisker: mean±s.e.m.. **E.** Cumulative variance explained across principal components for the full LP population, secondary optotagged cells, and non-secondary optotagged cells. **F.** Variance explained by the first three PCs. Bar and whisker: Mean±95% confidence interval (CI) of bootstrap iterations (subsampled to 30 cells). Statistical significance was measured using the CI of the difference between bootstrap distributions. No significant differences were detected (all p>0.25).

18

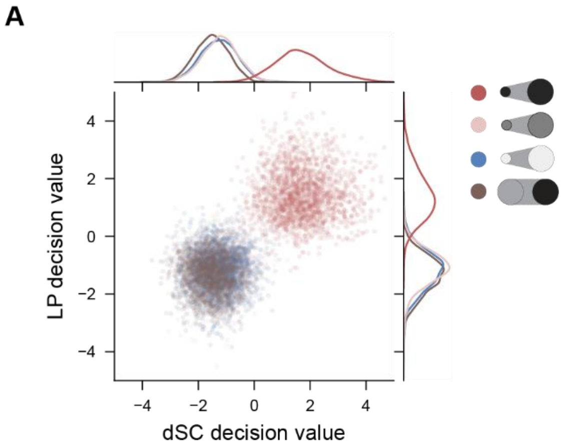


**Figure S7. Empirical distributions of threat decoder decision values. Related to Figure 8. A.** 2D state space defined by threat decoder decision values from *in vivo* experimental recordings in the dSC and LP (n = 24 dSC, 12 LP mice). Scatter points represent single trial decision values generated by randomly pairing bootstrapped trials across regions. Top and right curves show the probability density distributions for dSC and LP. Stimulus conditions: threat stimulus (black looming) and non-threat stimuli (gray looming, white looming, and dimming).

19

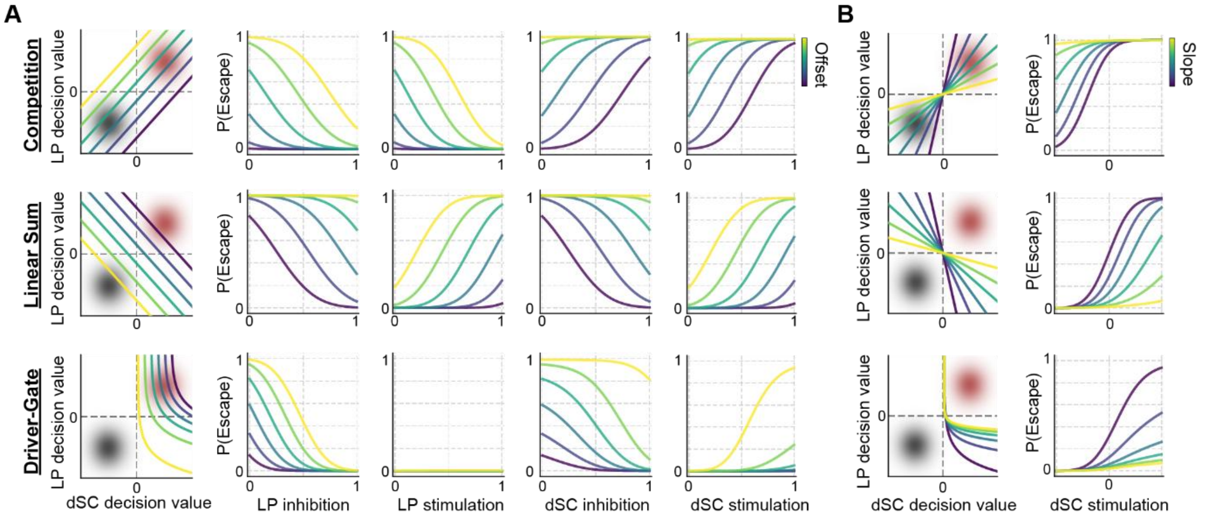


**Figure S8. Impact of decision boundary offsets and dSC stimulation. Related to Figure 8. A.** Decision boundaries across a range of offsets (leftmost column) and simulated escape probability across increasing strengths of *in silico* LP and dSC perturbations. **B.** Decision boundaries across a range of slopes (left) and simulated escape probability across increasing strengths of dSC stimulation.
